# Intact reading ability in spite of a spatially distributed visual word form ‘area’ in an individual born without the left superior temporal lobe

**DOI:** 10.1101/2021.09.15.460550

**Authors:** Jin Li, Hope Kean, Evelina Fedorenko, Zeynep Saygin

## Abstract

The visual word form area (VWFA) is an experience-dependent region in the left ventral temporal cortex (VTC) of literate adults that responds selectively to visual words. Why does it emerge in this stereotyped location? Past research shows the VWFA is preferentially connected to the left-lateralized frontotemporal language network. However, it remains unclear whether the presence of a typical language network and its connections with VTC are critical for the VWFA’s emergence, and whether alternative functional architectures may support reading ability. We explored these questions in an individual (EG) born without the left superior temporal lobe but exhibiting normal reading ability. We recorded fMRI activation to visual words, objects, faces, and scrambled words in EG and neurotypical controls. We did not observe word selectivity either in EG’s right homotope of the VWFA (rVWFA)—the most expected location given that EG’s language network is right-lateralized—or in her spared left VWFA (lVWFA), despite typical face selectivity in both the right and left fusiform face area (rFFA, lFFA). We replicated these results across scanning sessions (5 years apart). Moreover, in contrast with the idea that the VWFA is simply part of the language network that responds to general linguistic information, no part of EG’s VTC showed selectivity to higher-level linguistic processing. Interestingly, multivariate pattern analyses revealed sets of voxels in EG’s rVWFA and lVWFA that showed 1) higher within- than between-category correlations for words (e.g., Words-Words>Words-Faces), and 2) higher within-category correlations for words than other categories (e.g., Words-Words>Faces-Faces). These results suggest that a typical left-hemisphere language network may be necessary for the emergence of focal word selectivity within the VTC, and that orthographic processing can be supported by a distributed neural code.

## Introduction

In the past two decades, numerous regions in the ventral temporal cortex (VTC) have been identified and characterized that respond selectively to different high-level visual categories (e.g., faces: Kanwisher et al., 1997; places: Epstein & Kanwisher, 1998; bodies: Downing et al., 2001; for review, see Kanwisher, 2010). What are the origins of these specialized regions? How do human brains develop this functional topography? Some have hypothesized that the functional organization of the VTC may be innate and related to the evolutionary importance of certain high-level visual categories. Indeed, face perception and recognition abilities appear to be heritable (Wilmer et al., 2010; Zhu et al., 2010), and 2-9 month-old infants already show face- and place-responsive areas within expected locations in the VTC (Deen et al., 2017; Kosakowski et al., 2022). A related hypothesis is that pre-existing biases for certain visual attributes (e.g., retinotopy, Hasson et al., 2002; Malach et al., 2002; rectilinearity, Nasr et al., 2014) or perceptual dimensions (e.g., real-world size and animacy, Konkle & Caramazza, 2013) may predispose a brain region to become selective for particular visual categories. However, evolutionary pressures cannot explain the existence of a brain region that specializes for orthography (Hannagan et al., 2015)—the visual word form area (VWFA). The VWFA is a small region in the left lateral VTC that shows strong selectivity for visual words and letter strings in literate individuals (e.g., L. Cohen et al., 2003; Baker et al., 2007; Dehaene & Cohen, 2011; Hamamé et al., 2013). Perhaps surprisingly—given its late emergence—the VWFA is located in approximately the same location across individuals and scripts (Baker et al., 2007). What sets word forms apart from other visual categories and why does the VWFA land in this stereotyped location?

One compelling possibility is that the specialization of category-selective regions in the VTC is constrained by their differential connectivity to the rest of the brain (the Connectivity Hypothesis; Mahon & Caramazza, 2011; Martin, 2006). Indeed, previous work showed that category-selective responses can be predicted from both structural and functional connectivity (Osher et al., 2015; Saygin et al., 2012); further, these distinct connectivity patterns may already exist at birth and may drive future functional specialization (e.g., newborns show functional connectivity differences between lateral VTC which houses the VWFA vs. medial VTC: Barttfeld et al., 2018). Thus, written words may be processed in a stereotyped region within the left VTC due to this region’s pre-existing connectivity with the left-lateralized language network (e.g., Behrmann & Plaut, 2013; Dehaene et al., 2015; Martin, 2006). This network consists of left lateral frontal and lateral temporal areas and selectively supports language comprehension and production (e.g., Fedorenko et al., 2011).

Consistent with this idea, a number of studies have reported both anatomical and functional connections between the VWFA and the language network in neurotypical adults. For example, compared to the adjacent FFA, the VWFA shows stronger anatomical connectivity to the left superior temporal, anterior temporal, and inferior frontal areas (perisylvian putative language regions) (Bouhali et al., 2014). Several candidate white matter fascicles may serve to connect the VWFA with frontal and superior temporal language cortex (Wandell et al., 2012; Yeatman & Feldman, 2013). Similarly, using resting-state functional connectivity, Stevens et al. (2017) found that the individually defined VWFA connects to the posterior left inferior frontal gyrus as well as the left planum temporale, both part of the distributed left-hemisphere language network.

Moreover, a longitudinal study in children showed that the location of the VWFA could be successfully predicted by its connectivity patterns at the pre-reading stage (before any word selectivity within the VTC is observed), and that the most predictive connections of the relevant area of the VTC were with the frontal and temporal language areas (Saygin et al., 2016). Even more strikingly, this pattern of preferential connectivity appears to be already present in neonates (Li et al., 2020). These connections between the VWFA and the frontotemporal language network appear to be functionally important such that their damage or stimulation leads to reading difficulties. For example, a case study of a child with a missing arcuate fasciculus (AF, presumably connecting the VTC and other parts of the temporal cortex to parietal and frontal areas; Wandell et al., 2012) found impaired reading ability (Rauschecker et al., 2009). Similarly, a lesion in the white matter ventral to the angular gyrus resulted in alexia without agraphia, presumably disrupting the VWFA’s connections with the lateral temporal language areas (Greenblatt, 1976).

A different way to assess the importance of a language network in developing visual word selectivity is to ask whether language regions and the VWFA occupy the same hemisphere. In the majority of individuals, the language regions and the VWFA co-lateralize to the left hemisphere (LH) (e.g., Cai et al., 2010; Gerrits et al., 2019). In rare instances where neurotypical individuals show right-hemispheric (RH) language dominance, VTC activation during reading tasks also tends to be right-lateralized (e.g., Cai et al., 2008; Van der Haegen et al., 2012).

Another population where the language network is right-lateralized are individuals with congenital or early left hemisphere (LH) damage (e.g., Asaridou et al., 2020). In the presence of early LH damage, linguistic abilities tend to develop normally (e.g., Newport et al., 2017; Staudt, 2010; Staudt et al., 2001; see François et al., 2021 for a review). However, little is known about the effects of early LH damage on reading ability and on the neural architecture of visual word processing. In particular, if the left VTC is completely deafferentiated from the downstream LH language cortex at birth, does the VWFA emerge in the right VTC when the language network has no choice but to develop in the right hemisphere, or does it still emerge in the left VTC, due to some pre-existing bias (e.g., innate connectivity with any spared LH cortex)? Indeed, some previous studies show that language processing and reading *can* engage opposite hemispheres (e.g., Van der Haegen et al., 2012). Or—perhaps even more drastically—whatever hemisphere it emerges in, does the VWFA look atypical (e.g., less functionally selective or integrated into the language network, which in neurotypical individuals responds to linguistic demands across reading and listening modalities; e.g., Fedorenko et al., 2010; Vagharchakian et al., 2012; Regev et al., 2013)? Do these potential differences affect selectivity for other high-level visual categories in the VTC? And if the VWFA manifests atypically but the reading ability is normal, what does this tell us about the implementation-level flexibility with respect to orthographic processing? Here we investigate possible functional reorganization of the visual word selectivity in the absence of a typical left-lateralized language network. We have a unique opportunity to examine fMRI responses to stimuli from different visual categories in an individual (EG) born without the left superior temporal lobe (likely due to pre/perinatal stroke) but with lVTC largely intact. EG’s frontotemporal language network is completely remapped to the right hemisphere; no language-related responses, as assessed with fMRI, were observed in the remaining parts of EG’s left hemisphere (Tuckute et al., 2022). EG’s reading abilities (as well as other linguistic abilities) are intact. We here investigated a) whether, in the presence of a right-lateralized language network, a typical VWFA would emerge in the right VTC (in a typical location, or perhaps in other parts of the right VTC); b) whether any word selectivity is observed in the (spared) left VTC; and c) whether visual word processing could be taken over by brain regions that support general linguistic processing. To foreshadow the results, no word selectivity was observed in EG’s right or left VTC, despite typical selectivity for other visual categories; and brain regions that support high-level language processing did not distinguish between visual words and other visual categories, ruling out the possibility that univariate visual word processing is taking place within this high-level language network or within VTC. We then explored the possibility that orthographic processing is supported by still selective but more distributed neural populations using multivariate analyses, and indeed observed such ‘multivariate selectivity’ bilaterally, but manifesting more strongly in the right VTC.

## Methods

### Participants

#### Critical participant

The participant EG (fake initials; right-handed female with an advanced professional degree, 54 years old at the time of testing) contacted Dr. Fedorenko’s lab to participate in brain research studies. Based on her own report, the lack of the left superior temporal lobe (**Figure 2**) was discovered when she was 25 years old (in her first MRI scan in 1987) and being treated for depression. No known head traumas or injuries were reported as a child or adult. Several medical MRI scans were performed in subsequent years (1988, 1998, and 2013) and no change was observed compared to the initial scans. Importantly, EG did not report any difficulties in reading or general language abilities (see details below). She had also acquired fluency in a second language (Russian). EG was invited to participate in a series of behavioral and fMRI assessments at MIT. With respect to testing relevant to the current study, EG completed five runs of the VWFA localizer (see *The VWFA localizer task* section below) in October 2016 (session 1), and four runs of the same VWFA localizer in November 2021 (session 2). Our main analysis focused on session 1 (see *Data acquisition* section below), and we invited EG back for the second session to replicate results and also more critically, to search potential word-selective responses outside the VTC with whole-brain coverage (see Supplementary Methods: *Data acquisition for the VWFA localizer with a whole-brain coverage*). Written informed consent was obtained from EG, and the study was approved by MIT’s Committee on the Use of Humans as Experimental Subjects (COUHES).

#### Neurotypical controls

Twenty-five adults (11 female, mean age = 23.6 years old; age range 18- 38 years; standard deviation 5.21 years) from The Ohio State University (OSU) and the surrounding community were included in the present study. As part of ongoing projects exploring the relationship between brain function and connectivity, all participants completed a battery of fMRI tasks, including, critically, the same VWFA localizer task that EG completed (see *The VWFA localizer task* section below). All participants had normal or corrected-to-normal vision, and reported no neurological, neuropsychological, or developmental diagnoses. Written informed consent was obtained from all participants and the study was approved by Institutional Review Board at OSU. (It is worth noting that although the control group participants were younger than EG, further examination revealed no significant correlation between age and word selectivity (see *Definition of functional regions of interest and univariate analyses* section below) in our control group (r=0.199, p= 0.835; p-value was obtained by a permutation test)

#### Reading assessment (EG only)

To formally evaluate EG’s linguistic abilities, five standardized language assessment tasks were administered: i) an electronic version of the Peabody Picture Vocabulary Test (PPVT-IV) (Dunn & Dunn, 2007); ii) an electronic version of the Test for Reception of Grammar (TROG-2) (Bishop, 2003); iii) the Western Aphasia Battery-Revised (WAB-R) (Kertesz, 2006); iv) the reading and spelling components of PALPA (Kay et al., 1992); and v) an electronic version of the verbal components of the Kaufman Brief Intelligence Test (KBIT-2) (Kaufman & Kaufman, 2004). PPVT- IV and TROG-2 target receptive vocabulary and grammar, respectively. In these tasks, the participant is shown sets of four pictures accompanied by a word (PPVT-IV, 72 trials) or sentence (TROG-2, 80 trials) and has to choose the picture that corresponds to the word/sentence by clicking on it. WAB-R (Kertesz, 2006) is a more general language assessment developed for persons with aphasia. It consists of 9 subscales, assessing 1) spontaneous speech, 2) auditory verbal comprehension, 3) repetition, 4) naming and word finding, 5) reading, 6) writing, 7) apraxia, 8) construction, visuospatial, and calculation tasks, and 9) supplementary writing and reading tasks. Three composite scores (language, cortical, and aphasia quotients) were calculated from the subscales (the criterion cut-off score for diagnosis of aphasia is an aphasia quotient of 93.8). The verbal components of KBIT-2 include 1) the Verbal Knowledge subtest, which consists of 60 items measuring receptive vocabulary and general information about the world, and 2) the Riddles subtest consists of 48 items measuring verbal comprehension, reasoning, and vocabulary knowledge. Most relevant to the current investigation, the reading component of WAB-R includes comprehension of written sentences and reading commands; the supplementary reading tasks include reading of irregular words and nonwords. The reading component of PALPA (Kay et al., 1992) consists of 21 tasks (tasks #18 through 38), which focus on single letter recognition, visual lexical decision, out-loud reading (of words with diverse lexical properties and sentences), and homophone definition (see Supplementary Table 2 for details).

To further investigate any reading or visual word processing deficiencies which may not be apparent in the standard linguistic and reading measures reported above, we also had EG complete a speeded reading task and compared her performance to that in an independent large sample of neurotypical adults (see Ryskin et al., in preparation, for details). Briefly, twelve- word-long sentences were presented word-by-word at varying speeds. The original presentation speed was based on the natural out-loud reading speed (as recorded by a female native English speaker): each word was visually presented for the number of ms that it took the reader to say the word. The speed was manipulated by compressing the sentence presentations to 80%, 60%, 50%, 45%, 40%, 35% and 30% of the original speed (100%). Participants were asked to type in as many words as they were able to discern after each sentence, and the accuracy of participants’ responses (how many words of the sentence they typed in correctly, not taking into account minor typos) was recorded.

#### Data acquisition

EG’s data were collected on a 3 Tesla Siemens Trio scanner with a 32-channel head coil at the Athinoula A. Martinos Imaging Center at the McGovern Institute for Brain Research at MIT. Data of the control group participants were acquired on a 3 Tesla Siemens Prisma scanner with a 32-channel head coil at the Center for Cognitive and Behavioral Brain Imaging at OSU. To ensure that any differences between EG and our control group were not due to scanner differences, we compared word selectivity in the current control group to a smaller group of adults recruited at MIT and scanned with the same scanner and protocols as EG; no differences were observed (Supplementary Table 1). For both EG and controls, a whole-head, high resolution T1-weighted magnetization-prepared rapid acquisition with gradient echo (MPRAGE) scan was acquired (EG: repetition time (TR) = 2530 ms, echo time (TE) = 3.48 ms; voxel resolution = 1.0 mm^3^; the control group: TR = 1390 ms, TE = 4.62 ms, voxel resolution = 1.0 mm^3^). Functional images for the VWFA task were acquired with the same echo-planar imaging (EPI) sequence for both EG (session 1, the first visit) and controls: TR = 2000ms, TE = 30ms, 172 TRs, 100 × 100 base resolution, voxel resolution=2.0 mm^3^, field of view (FOV) = 200mm; 25 slices approximately parallel to the base of the temporal lobe to cover the entire ventral temporal cortex. Unless otherwise noted, results presented are from EG’s first visit where the acquisition parameters were identical for EG and controls. To search for potential word selectivity outside the VTC, we invited EG back recently and collected data from the same VWFA task with a slightly different protocol to get whole-brain coverage (see Supplementary Methods). EG also completed a language localizer task during session 1: EPI sequence with TR = 2000ms and TE = 30ms, 227 TRs, 96 × 96 base resolution, voxel resolution =2.1 × 2.1 × 4.4 mm3, FOV = 200mm, 31 near-axial slices acquired in the interleaved order.

#### The VWFA localizer task

A VWFA localizer was used to define high-level category-selective regions and to measure category-selective responses (see Saygin et al., 2016, for details). Briefly, black and white line drawings of words, scrambled words, objects, and faces, along with the fixation condition were shown in a blocked design. A grid was overlaid on top of the stimuli so that all stimulus types (not just scrambled words) had edges. Each stimulus was presented for 500ms (ISI=0.193s) and overlaid on a different single-color background, and 26 stimuli (including 2 repetitions) were presented in each block. Each run consisted of 19 blocks (4 blocks per condition and 3 fixation blocks), and participants performed a one-back task. The stimuli are available for download at http://www.zeynepsaygin.com/ZlabResources.html. EG completed 5 runs, and participants in the control group completed 2 runs. Note that previous studies using the same task indicated that 2 runs of data are sufficient to successfully identify the VWFA in a neurotypical population (Saygin et al., 2016); here, we acquired more runs for EG to ensure that we had sufficient power and that the results obtained for EG were stable across runs (see *Definition of functional regions of interest and univariate analyses* section for details).

#### The language localizer task

During session 1, EG also completed a language localizer that was adapted from Fedorenko et al. (2010). Briefly, there were four types of stimuli: English sentences, scrambled sentences (i.e., word lists), jabberwocky sentences (sentences where all the content words are replaced by pronounceable nonwords, like, for example, “florped” or “blay”), and nonword sequences (scrambled jabberwocky sentences) (see Fedorenko et al., 2010 for details). Stimuli were presented in both visual and auditory blocks, each lasting 22s and 26s respectively. Each run consisted of 20 blocks (4 blocks per stimulus type × 2 modalities, and 4 fixation blocks), and the order was counterbalanced across runs. Note in visual blocks, stimuli were presented sequentially, that is, one word/nonword at a time. To control attention demands, a probe word/nonword was presented at the end of each trial, and subjects had to decide whether the probe appeared in the immediately preceding stimulus. The English sentences (En) and Nonword sequences (Ns) were the critical conditions to identify language-selective responses (i.e., high-level linguistic information: lexico-semantic and syntactic).

#### Preprocessing and fMRI analysis

Data were analyzed with Freesurfer v.6.0.0 (http://surfer.nmr.mgh. harvard.edu/), FsFast (https://surfer.nmr.mgh.harvard.edu/fswiki/FsFast), FSL (https://fsl.fmrib.ox.ac.uk/fsl/fslwiki/FLIRT) and custom MatLab code. All structural MRI data were processed using a semiautomated processing stream with default parameters (recon-all function in Freesurfer: https://surfer.nmr.mgh.harvard.edu/fswiki/recon-all/), which includes the following major steps: intensity correction, skull strip, surface co-registration, spatial smoothing, white matter and subcortical segmentation, and cortical parcellation. Cortical gray matter and ventral temporal cortex masks were created based on the Desikan-Killiany (Desikan et al., 2006) parcellation in native anatomy for each subject. The success of cortical reconstruction and segmentation was visually inspected for EG.

Functional images were motion-corrected (time points where the difference in total vector motion from the previous time point exceeded 1mm were excluded), data from each run were registered to each individual’s anatomical brain image using bbregister, and resampled to 1.0x1.0x1.0mm^3^. For EG, instead of registering the functional image of each run to the anatomical brain separately, we aligned the functional images of the first four runs to the last run (which had successful functional-anatomical cross-modal registration with bbregister) with linear affine transformation (FLIRT); then the functional-anatomical transformation for the last run was applied to all functional runs and was visually inspected (tkregisterfv) for functional to anatomical alignment.

The preprocessed functional data were then entered into a first-level analysis. Specifically, data were detrended, smoothed (3mm FWHM kernel), and the regressor for each experimental condition (Words, Scrambled Words, Objects, and Faces) was defined as a boxcar function (i.e., events on/off) convolved with a canonical hemodynamic response function (a standard gamma function (d = 2.25 and t = 1.25)). Orthogonalized motion measures from the preprocessing stage were used as nuisance regressors for the GLM. Resulting beta estimates for each condition and contrasts of interest between categories (i.e., Words > Others categories and Faces > Others categories) were used in further analyses. For multivariate analyses, no spatial smoothing was applied.

#### Definition of functional regions of interest and univariate analyses

The subject-specific functional regions of interests (fROIs) were defined with a group-constrained subject-specific (GcSS) approach (Fedorenko et al., 2010). In this approach, individual fROIs are defined by intersecting participant-specific fMRI contrast maps for the contrast of interest (e.g., Words > Other conditions) with some spatial constraint(s) or ‘parcel(s)’, which denote the area(s) in the brain within which most individuals show responses to the relevant contrast (based on past studies with large numbers of participants). In the present study, the VWFA and the FFA parcels were derived from previous studies (FFA, Julian et al., 2012; VWFA, Saygin et al., 2016) (**Figure 2**) and were generated based on probabilistic maps of functional activation for the relevant contrasts in independent groups of participants. We registered these parcels to our participants’ own anatomy using the combined volume and surface-based (CVS) non-linear registration method (mri_cvs_register; Postelnicu et al., 2009). After mapping the functional parcels to each participant’s brain, we defined the VWFA fROI by selecting within the VWFA parcel the top 10% of most active voxels for the contrast Words > Other conditions (i.e., scrambled words, objects, and faces). Similarly, we defined the FFA by selecting within the FFA parcel the top 10% of most active voxels for the contrast Faces > Other conditions. For the control participants, we used run 1 to define the fROIs, and the run 2 to extract percent signal changes (PSCs; beta estimates divided by baseline activation) for each of four experimental conditions; the same procedure was repeated with run 2 to define the fROIs and run 1 to extract PSCs, and the average result for each subject was used in further analyses.

For EG, who had 5 runs of data, this procedure was performed iteratively for every 2-run combination (e.g., defining fROIs with run 1 and extracting PSCs from run 2, then defining with run 2 and extract from run 1, and results were averaged; the same procedure were repeated for all 2-run combinations (e.g., 1&3, 1&4, etc.; 10 combinations in total). When comparing EG to controls, results from ten run combinations were averaged to derive a single estimate per condition per fROI. Note that although the parcels are relatively large, by design (in order to accommodate inter-individual variation in the precise location of the functional region), and can overlap, the resulting fROIs within an individual are small and do not overlap (Saygin et al., 2016). We further calculated the selectivity indices for words and faces with the following formula: (PSC to the condition of interest – average PSC of the remaining conditions)/(summed PSC for all four conditions); note that when calculating selectivity, we adjusted for baseline activation following previous studies (Simmons et al., 2007; Szwed et al., 2011) in order to correct for potential bias induced by negative activation that are sometimes observed in fMRI studies.

#### Multivariate analyses: Split-half correlations

To further examine whether visual words may be represented and processed in a spatially distributed manner, we performed a multivariate pattern analysis (MVPA) to measure distinctive activation patterns for different conditions. The analyses were performed with CoSMoMVPA toolbox (Oosterhof et al., 2016) (https://w.cosmomvpa.org/). In line with the approach introduced in Haxby et al. (2001), we examined split-half within-category and between-category correlations. In particular, a searchlight (radius = 3 voxels) was created for each voxel within the VTC, and response patterns (i.e., beta estimates for each of the four conditions, normalized by subtracting mean responses across all conditions) were extracted from each searchlight.

Before performing the critical analysis, we asked whether the overall multivariate representation structure of the VTC is typical in EG. To do so, we constructed a representational similarity matrix (RSM) from pairwise similarities (i.e., correlations) based on the voxel-wise response patterns in a given searchlight area (defined above) to different conditions (e.g., the correlation between activation patterns across voxels to Words and Faces in a given searchlight area).

Correlations of all searchlights within the VTC were then Fisher z-transformed and averaged. This resulted in one 4x4 RSM from two runs of data for each participant in the control group; for EG, RSMs from ten run combinations were averaged to get a single RSM. We then calculated RSM similarity (correlation) between EG and controls, and tested whether this correlation was different from the correlations between any given control individual and the rest of the control group (within-controls correlations; similarity of RSM of each subject to the average RSM of the remaining control subjects).

Then, similarity of response patterns within a category (e.g., Words-Words or Faces-Faces) vs. between categories (e.g., Words-Faces) was calculated within the VWFA and FFA parcel boundaries by Pearson’s correlation in a pair-wise manner, and then Fisher z-transformed. We identified voxels of interest that satisfied the following two criteria: 1) voxels whose local neighborhoods showed higher within- than between-category correlations (and which therefore represent categories distinctively); and 2) voxels whose local neighborhoods showed higher within-condition correlations for a particular category (e.g., Words-Words) than within-condition correlations for other categories (e.g., Faces-Faces). The second criterion identified voxels which represent a particular category (e.g., visual words) in a more selective fashion. We refer to such voxels as multivariate-selective voxels.

To examine hemispheric differences, we computed the number of voxels that exhibited multivariate selectivity for words or faces in the two hemispheres (within the relevant parcels). To control for the difference in the size of the search spaces (i.e., the parcels), we divided the number of the multivariate-selective voxels by the total size of the relevant parcel.

#### Statistical analyses

Paired t-tests were used for comparisons between conditions for EG (across ten run split combinations) and within the control group. For all analyses where we compared EG’s response to the control group, we used a Crawford-Howell modified t-test (Crawford & Howell, 1998), which is widely used in single-case studies because it accounts for the variance of the control group, and the percentage of false positives remains the same for various sizes of the control sample (N=5-50) (Crawford et al., 2009). This frequentist approach provided the point estimate (p-value) for the proportion of the control population that will obtain a score more extreme than the critical participant’s score. One-tailed tests were reported for the critical (directional) hypothesis tested in the current study (unless noted otherwise). In addition, we computed the Bayesian 95% credible interval (a Bayesian alternative to the modified t-test; Crawford & Garthwaite, 2007) to demonstrate the range of p-values based on 10,000 iterations. To assess the significance of RSM correlations between EG and controls, we generated a null distribution of correlation values by shuffling the matrix of EG and controls (i.e., randomizing the labels of values in the RSMs) and then correlating the new shuffled matrices. This procedure was repeated 10,000 times to create the null distribution of the correlation values. The p-value was calculated by counting the number of correlations in the null-distribution that were higher than the correlation value based on the correct category labels, and then divided by 10,000.

## Results

### Does EG have normal reading ability?

In line with her self-report, EG performed within normal range on all language assessment tasks. Her accuracy was 90% on PPVT, 99% on TROG-2, and she obtained the scores of 97.6, 98.6, and 98.4 on the aphasia, language, and cortical quotients of the WAB-R respectively, and a score of 130 (98th percentile) on the KBIT-verbal. EG’s performance was therefore not distinguishable from the performance of neurotypical controls. With respect to the reading assessments, EG made no errors on the main reading component of WAB-R, no errors in the reading of irregular words, and one error in the reading of nonwords (reading ‘gobter’ instead of ‘globter’). For the PALPA tasks, she made no errors on tasks that focus on single letters (tasks #18-23), no errors on the visual decision tasks (tasks #24-27), no errors on the out-loud reading tasks (tasks #29-37), and no errors on the homophone definition task (task #38). For the homophone decision task (task #28), EG made three errors (out of the 60 trials; all were made on nonword pairs: she did not judge the following pairs as sounding the same: heem-heam, byme-bime, and phex-feks).

This performance is on par with neurotypical controls.

Consistent with the results from these standard reading measures, behavioral performance during the 1-back VWFA localizer task in the scanner also showed that EG’s response accuracy and response time were not different from controls for all visual categories (**Figure 1A**, Supplementary Table 1a). Critically, EG’s performance on the speeded reading task was also within the range of the control distribution (see Methods for details) even for the fastest presentation rates (**Figure 1B**; Supplementary Table 1b). This result demonstrates that not only does EG perform within the typical range on temporally unconstrained/self-paced reading assessments, but her reading mechanisms are not compromised in terms of their speed.

**Figure 1.**
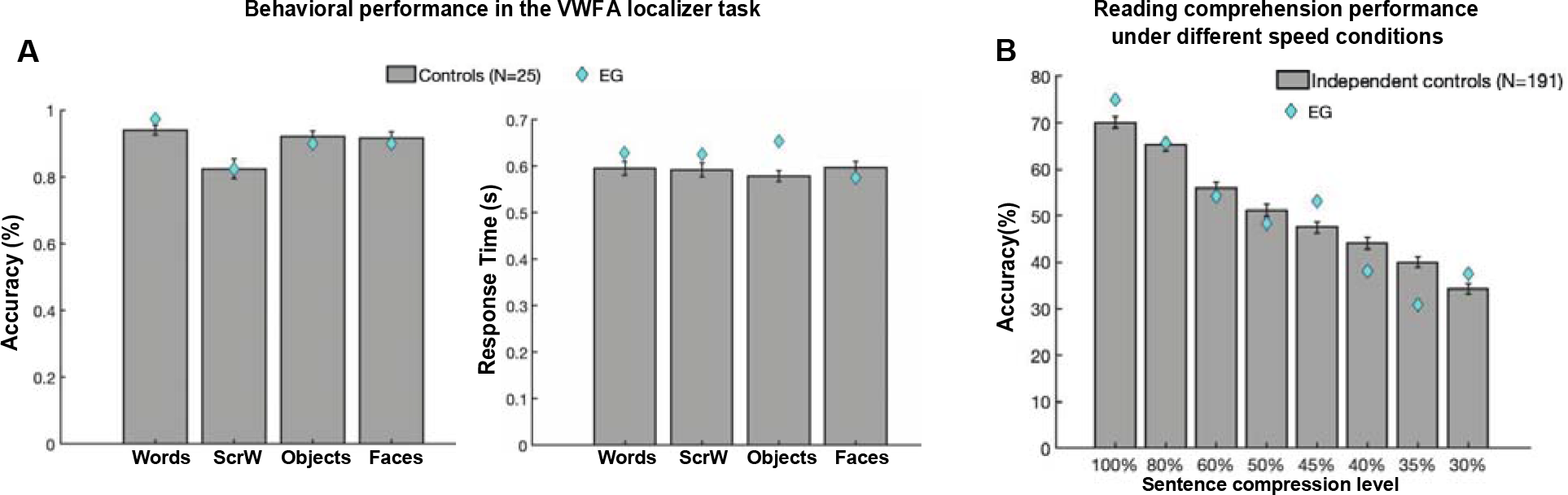
Comparing EG’s reading performance with neurotypical controls. (A) Accuracy (left) and response time (right) during the 1-back VWFA localizer task. (B) Proportion of words typed that matched the words in the target sentence in different speed conditions.

**Figure 2.**
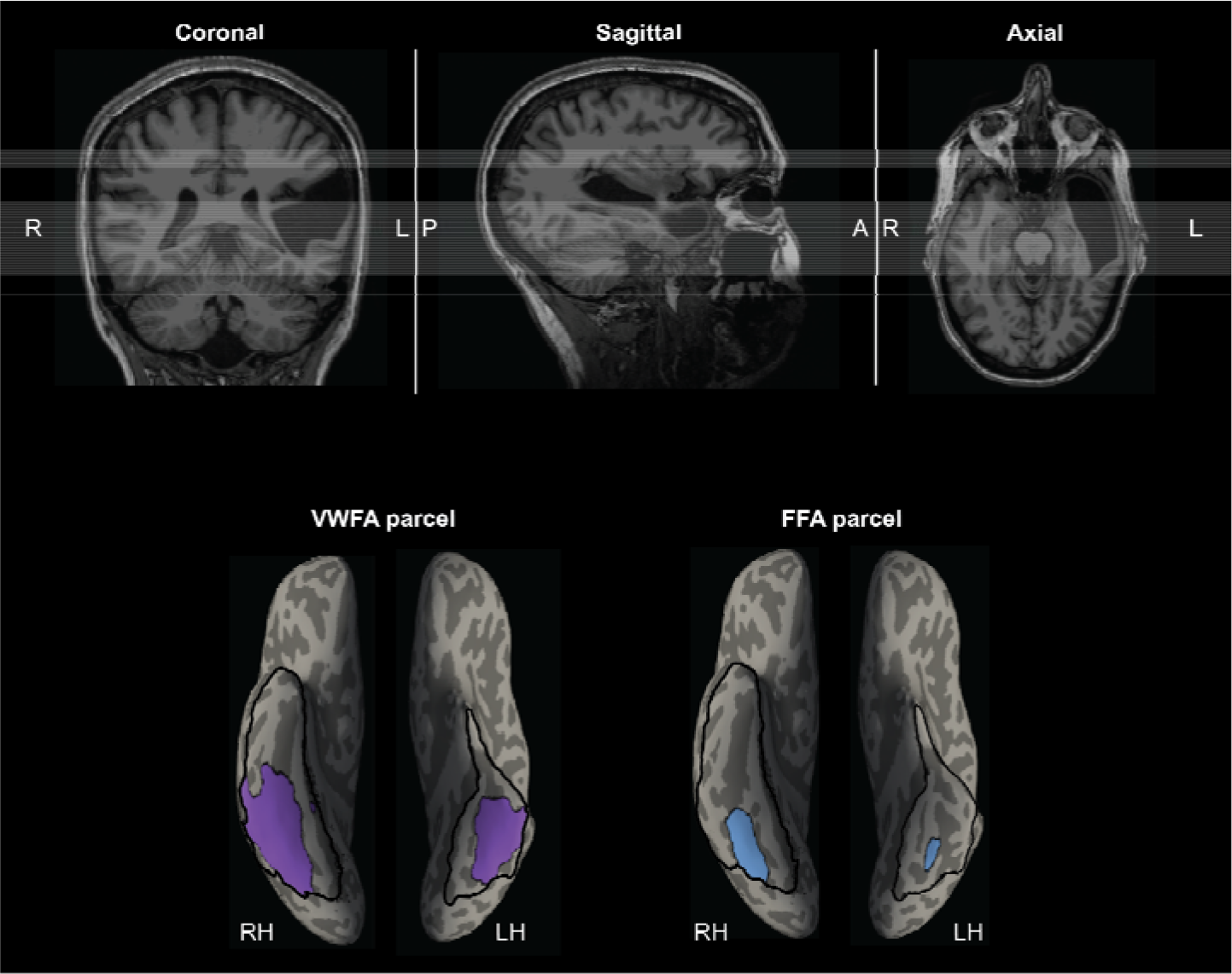
EG’s MRI showing the missing cortex and the parcels for the right and left VWFA and FFA. Top, T1-weighted images in coronal, sagittal, and axial views. Bottom, the VWFA (purple) and FFA (blue) parcels are projected on EG’s reconstructed surface. The parcels served as spatial constraints in defining the fROIs (see Methods), but we also explored the entire VTC for category selectivity (outlined with black solid lines). By design, the parcels are relatively large (to accommodate inter-individual variability in the precise locations of these areas) and therefore can overlap, but the individual fROIs are small and do not overlap. Note that even though part of the anterior lVTC is missing in EG, the stereotypical locations for both the VWFA and FFA are spared.

Altogether, EG appears to have intact linguistic, reading, and visual word recognition ability.

### Is word selectivity observed in the right hemisphere when the language network is located in the right hemisphere from birth due to early left-hemisphere damage?

After confirming normal reading ability in EG, we moved on to our main analysis to examine univariate word selectivity in EG’s rVWFA. With the VWFA localizer, we defined the individual-specific rVWFA fROIs, in EG and controls, within a spatial constraint (rVWFA parcel; see **Figure 2** and Methods) by contrasting Words versus all other categories (Words > Others). We then extracted the activation to all four conditions from independent data (see Methods for details). In neurotypical literate individuals, word selectivity is strongly left-lateralized (McCandliss et al., 2003); selective responses to visual words in the right homotope of lVWFA are less frequently observed and are less spatially consistent across individuals.

Results from our control group are in line with this picture: we found no selective activation to Words as compared to other visual categories in the rVWFA (Words vs. Scrambled Words: t(24)=-0.548, p= 0.588; Words vs. Objects: t(24)=467, p= 0.644; Words vs. Faces: t(24)=-0.052, p= 0.959; **Figure 3A**).

**Figure 3.**
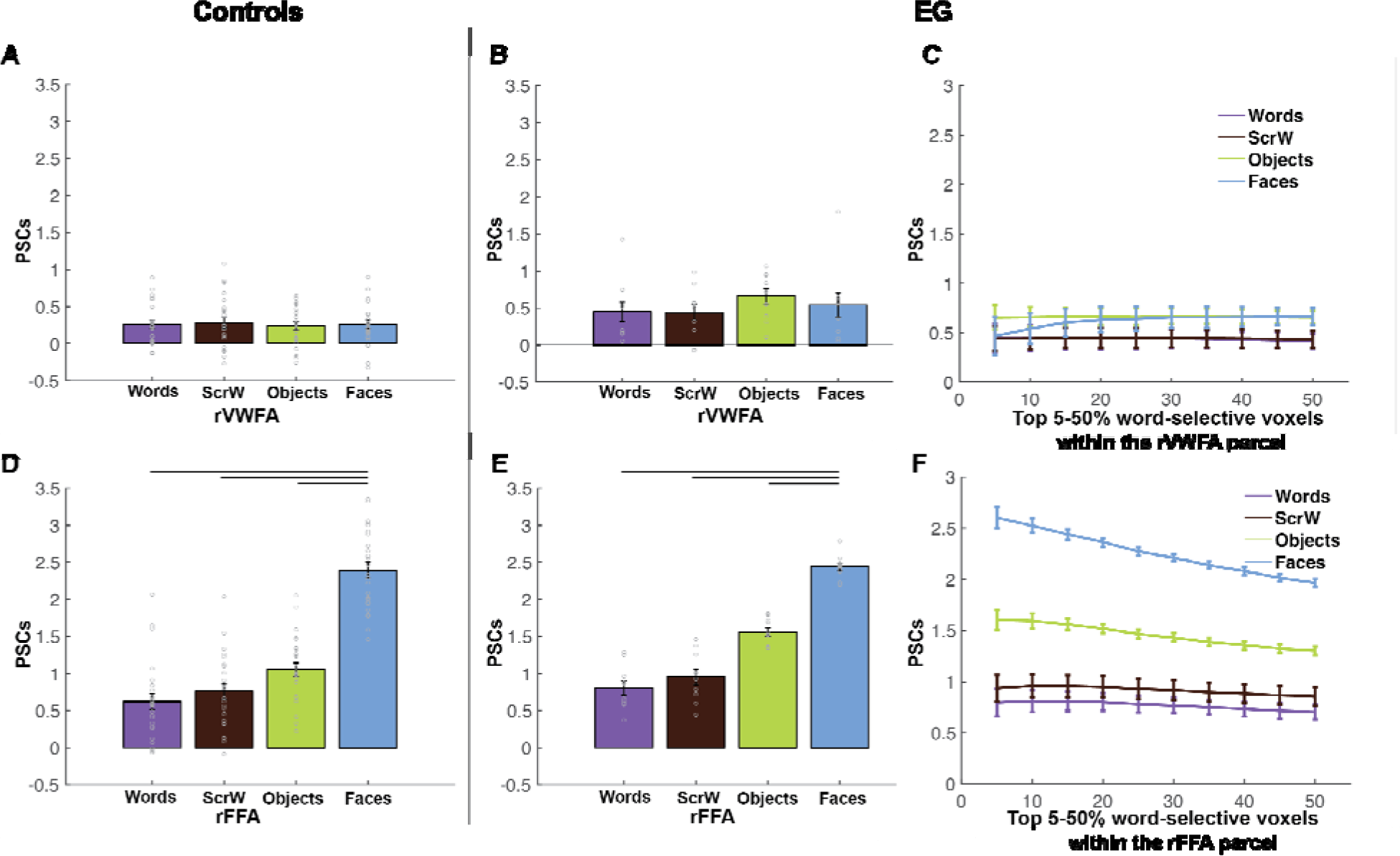
Responses to four conditions in the rVWFA and rFFA for EG and the control group. (A) Bar plots show mean PSCs to the four conditions estimated in independent data within individually defined rVWFA fROIs (i.e., top 10% word-selective voxels within the rVWFA parcel) for the control group. (B) Mean PSCs to the four conditions estimated in independent data within the individually defined rVWFA fROI for EG. Here and in E, the results are averaged across run combinations. (C) Parametrically decreasing the threshold for defining the rVWFA fROI from the top 5% to 50% word-selective voxels within the rVWFA parcel. Number of voxels in the rVWFA fROIs: 5%=384 voxels, 50%=3007 voxels. Here and in F, average PSCs across run combinations are shown for each threshold. (D) Mean PSCs to the four conditions estimated in independent data within individually defined rFFA fROIs for the control group. (E) Mean PSCs to the four conditions estimated in independent data within the individually defined rFFA fROI for EG. (F) Parametrically decreasing the threshold for defining the rFFA fROI from the top 5% to 50% face-selective voxels within the rFFA parcel. Number of voxels in the defined rFFA fROIs: 5%=90 voxels, 50%=699 voxels. In the bar plots, dots correspond to individual data points for each condition (controls: n=25 participants; EG: n=10 run combinations, from ten iterations). Horizontal bars reflect significant paired t-tests p < 0.05. Error bars in both the bar and line plots denote standard errors of the mean by participants (for the control group) and by run combinations (for EG).

Critically, in EG, whose language network is located in the right hemisphere, with no language responses anywhere in the left hemisphere (Tuckute et al., 2022), we asked whether word selectivity is also observed in the right hemisphere. Surprisingly, no word selectivity was observed in EG’s rVWFA: activation to Words in her rVWFA fROI did not significantly differ from other categories (Words vs. Scrambled Words: t(9)=0.056, p= 0.956; Words vs. Faces: t(9)=-1.107, p= 0.297; Words vs. Objects: t(9)=-1.661, p= 0.131) (**Figure 3B**). To ensure that this result is not due to the choice of a particular threshold (i.e., top 10%) that we used to define the rVWFA fROI, we performed the same analysis as above at a range of thresholds (5%-50%). The lack of word selectivity was stable across thresholds (**Figure 3C**). We then compared EG to the controls, and found that there was a trend whereby responses to Words (relative to baseline) were slightly higher in EG compared to the control group (t(24)=0.691, p= 0.248, modified t-test; 95% Bayesian CI [0.116, 0.382]), but this difference was not significant.

To test whether selectivity for other visual categories in the right VTC is typical in EG, we examined face selectivity in EG’s rFFA, which is spatially proximal to the rVWFA. Similar to the rVWFA analysis above, we defined the rFFA fROI by contrasting Faces versus other categories (Faces > Others), and extracted the activation to the four conditions from independent data (see Methods for details). EG’s face selectivity remained intact (**Figure 3E)** and did not differ from that of the control group (**Figure 3D**; t(24)=0.083, p= 0.467, modified t-test, Bayesian 95% CI [0.337 0.500]). EG’s rFFA fROI showed significantly higher responses to Faces than to other conditions (Faces vs. Words: t(9)=20.816, p= 6.380×10^-9^; Faces vs. Scrambled Words: t(9)=12.390, p= 5.862×10^-7^; Faces vs. Objects: t(9)=17.723, p=2.629×10^-8^; **Figure 3E)**, just like what we observed in the control group (Faces vs. Words: t(24)=16.466, p= 1.403×10^-14^; Faces vs. Scrambled Words: t(24)=14.895, p= 1.265×10^-13^; Faces vs. Objects: t(24)=13.837, p=6.213×10^-13^; **Figure 3D**). Moreover, the selective face responses in EG’s rFFA was observed across all thresholds used to define the rFFA (**Figure 3F**). We further calculated the strength of selectivity (selectivity index; Simmons et al., 2007) for both Words and Faces within rVWFA and rFFA respectively, by taking the difference between the condition of interests and the rest conditions and divided by the sum of all conditions (see Methods for details). We found that, consistent with previous observations, controls showed no word selectivity (i.e., compared to zero) in the rVWFA (t(24)= 0.157, p= 0.876) but significant face selectivity in rFFA (t(24)= 10.422, p= 2.178×10^-10^). Importantly, EG’s selectivity was not different from controls for both Words and Faces (Words: t(24)= -0.285, p= 0.389, modified t-test, Bayesian 95% CI [0.265 0.500]; Faces: t(24)= -0.718, p= 0.240, modified t-test, Bayesian 95% CI of EG’s activation [0.117 0.376]), again indicating the lack of word selectivity in the rVWFA and normal face selectivity in the rFFA in EG.

To ensure that we did not miss any possible word-selective voxels by applying a predefined spatial constraint (i.e., VWFA parcel) and to account for the possibility that EG’s VWFA may be located in a different part of the visual cortex, we searched for word selectivity within the entire rVTC mask for EG. Specifically, different thresholds from top 1% to top 10% were used to define the most word-selective voxels (Words > Others). Even within this broad mask, no word-selective responses were observed in independent data across all thresholds; in fact, the responses were lowest to words than the other three conditions (**Figure 4A**). In contrast, robust face-selective responses were observed in independent data across all thresholds when searching for face-selective voxels (Faces > Others) (**Figure 4B**). We also performed an analysis where we restricted the search space only to rVTC voxels that significantly and consistently responded to visual stimuli. Similar results were observed (Supplementary Figure 1C) where no word selectivity was observed in EG’s right visual cortex.

**Figure 4.**
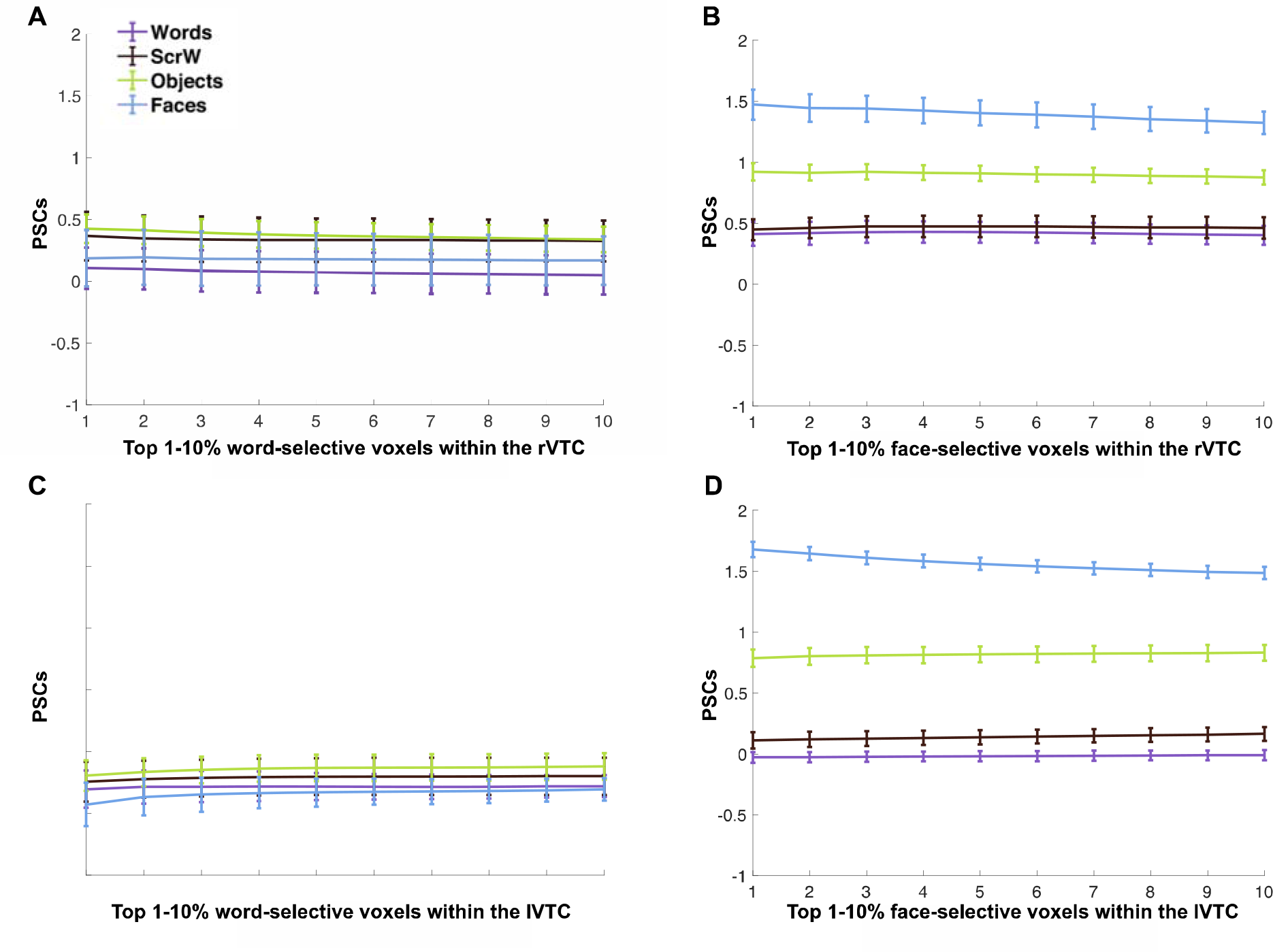
Mean PSCs in the rVTC and lVTC at different thresholds for EG. (A-B) Parametrically decreasing the threshold for defining word-selective (Words > Others) and face-selective (Faces > Others) voxels from the top 1% to 10% within the rVTC. Mean PSCs across run combinations (from 10 iterations) are shown for each threshold. (C-D) Parametrically decreasing the threshold for defining word-selective and face-selective voxels from the top 1% to 10% within the lVTC. Mean PSCs across run combinations are shown for each threshold. Number of selected voxels: rVTC: 1%=216 voxels, 10%=2164 voxels; lVTC: 1%=154 voxels, 10%=1536 voxels.

Moreover, we supplemented our main analyses, which rely on the Words > Others contrast to define the VWFA, as is commonly done in the literature (e.g., Dehaene-Lambertz et al., 2018; Rosenke et al., 2021), with another analysis that relies on a less stringent localizer contrast: Words vs. Scrambled Words (Glezer et al., 2009; Lerma-Usabiaga et al., 2018). Unlike the Words > Others contrast, this contrast does not control for semantics or visual stimulus complexity. Even with this broader contrast, we found no word selectivity within EG’s rVTC across thresholds (Supplementary Figure 1A).

Finally, some have claimed that even in neurotypical individuals responses to visual words in the VTC do not reflect orthographic processing (Price & Devlin, 2003) but that, instead, the VWFA is part of the language network (Price & Devlin 2011) and thus should show selectivity for linguistic stimuli in general (not just visually presented ones). Note that this hypothesis is not consistent with much existing evidence (e.g., Dehaene et al., 2004; Dehaene & Cohen, 2011) including data from neurotypical controls presented here (see *Is there any word selectivity in the spared left VTC* section below). But perhaps for populations with differently organized brains, this hypothesis has merit. For example, Kim et al. (2017) found that in congenitally blind individuals, whose visual cortex has long been known to show reorganization (e.g., Röder et al., 2002; Lane et al., 2015; Striem-Amit & Amedi, 2014), the anatomical location of the (putative) VWFA responded to both Braille and to a grammatical complexity manipulation for auditorily presented sentences. As a result, we considered the possibility that while EG lacks a right-lateralized VWFA that is selective to visual words over other visual categories, perhaps a part of her right VTC would respond to linguistic stimuli in general (either visually or auditorily presented) and that maybe this putative language region in the VTC supports EG’s normal reading ability. To explore this possibility, we implemented two analyses. First, we examined the same fROI as above (using the visual Words > Others contrast to define the rVTC fROI at different thresholds) to see if it exhibits language selectivity (as defined by higher responses to meaningful English sentences (En) than to sequences of Nonwords (Ns) (English sentences > Nonword sequences) presented either auditorily or visually (see Methods). We found no preferential activation to high-level linguistic information (i.e. no significant Sentences > Nonwords effect for sentences presented either auditorily or visually; Fedorenko et al., 2010) in the rVTC (Supplementary Figure 2A). Further, neural responses to the four visual categories (from the VWFA localizer) were not distinguishable from those for the conditions of the auditory language task (either meaningful English sentences or nonword sequences), suggesting that EG has no univariate response selectivity in rVTC to either visual words or linguistic stimuli in general (i.e. rVTC fROI defined by Words> Others does not show selectivity to words, visual stimuli in general, or either visual or auditory language; Supplementary Figure 2A). In contrast, the face-selective rVTC fROI showed consistently higher activation to faces than to all conditions in the language (and VWFA) localizer across all thresholds (Supplementary Figure 2B), suggesting that this result was specific to word selectivity and not visual category-selective areas in general. Second, we explored the possibility that there exists an amodal language region (e.g., semantics) somewhere in the VTC, outside the boundaries of the VWFA parcel, probably more anterior than the canonical location of the VWFA based on previous observations (e.g., Mummery et al., 2000; Schwartz et al., 2009); if so, perhaps this region would show selectivity to visual words as compared to other visual categories (note that this analysis is more of a reality check because such a region would have been picked out in the analysis searching for word selectivity across the VTC). Interestingly, we did not find a language-selective rVTC fROI that showed consistently higher activation to Sentences than to Nonword sequences (visually or auditorily presented) (Supplementary Figure 2C) even in the anterior part of the rVTC in EG. So, although EG’s language network was right-lateralized and showed canonical selectivity to linguistic stimuli (Tuckute et al., 2022), we found no evidence of a right VTC region that showed visual word selectivity or high-level linguistic selectivity.

### Is there any word selectivity in the spared left VTC?

Because we did not observe a right-lateralized VWFA and because the left VTC was largely intact, we also asked whether a canonical lVWFA may have developed in EG (perhaps due to some specific visual features of word forms that are better represented in the left hemisphere/right visual field (e.g., Hsiao & Lam, 2013; Seghier & Price, 2011; Tadros et al., 2013). In the control group, we observed robust word selectivity in the lVWFA (as expected from previous studies), with Words eliciting greater activation than each of the other conditions: Words vs. Scrambled Words: t(24)=6.733, p=5.779×10^-7^; Words vs. Objects: t(24)=4.983, p= 4.345×10^-5^; Words vs. Faces: t(24)=6.425, p= 1.211×10^-6^; **Figure 5A**). In contrast, we found no word selectivity in EG’s lVWFA (**Figure 5B**): activation to Words was around baseline and lower than the response to other categories, although the differences did not reach significance (Words vs. Scrambled Words: t(9)=-1.509, p= 0.165; Words vs. Objects: t(9)= -1.192, p= 0.264; Words vs. Faces: t(9)=-1.493, p= 0.170). Moreover, EG’s activation to Words was significantly lower than the control group’s (t(24)=-2.299, p= 0.015, modified t-test; 95% Bayesian CI [3.994×10^-5^, 0.046]). This result was stable across thresholds that were used to define the lVWFA fROI (**Figure 5C**), and we did not observe word selectivity in EG when we searched across the entire lVTC with varying thresholds (**Figure 5C**), visually-responsive voxels in the lVTC (Supplementary Figure 1D), or when we used a less stringent contrast (Words vs. Scrambled Words; Supplementary Figure 1B). Finally, we performed the same analysis as above where we explored linguistic selectivity using a language localizer, and found no evidence of a language-selective fROI within the lVTC (Supplementary Figure 2D-F).

**Figure 5.**
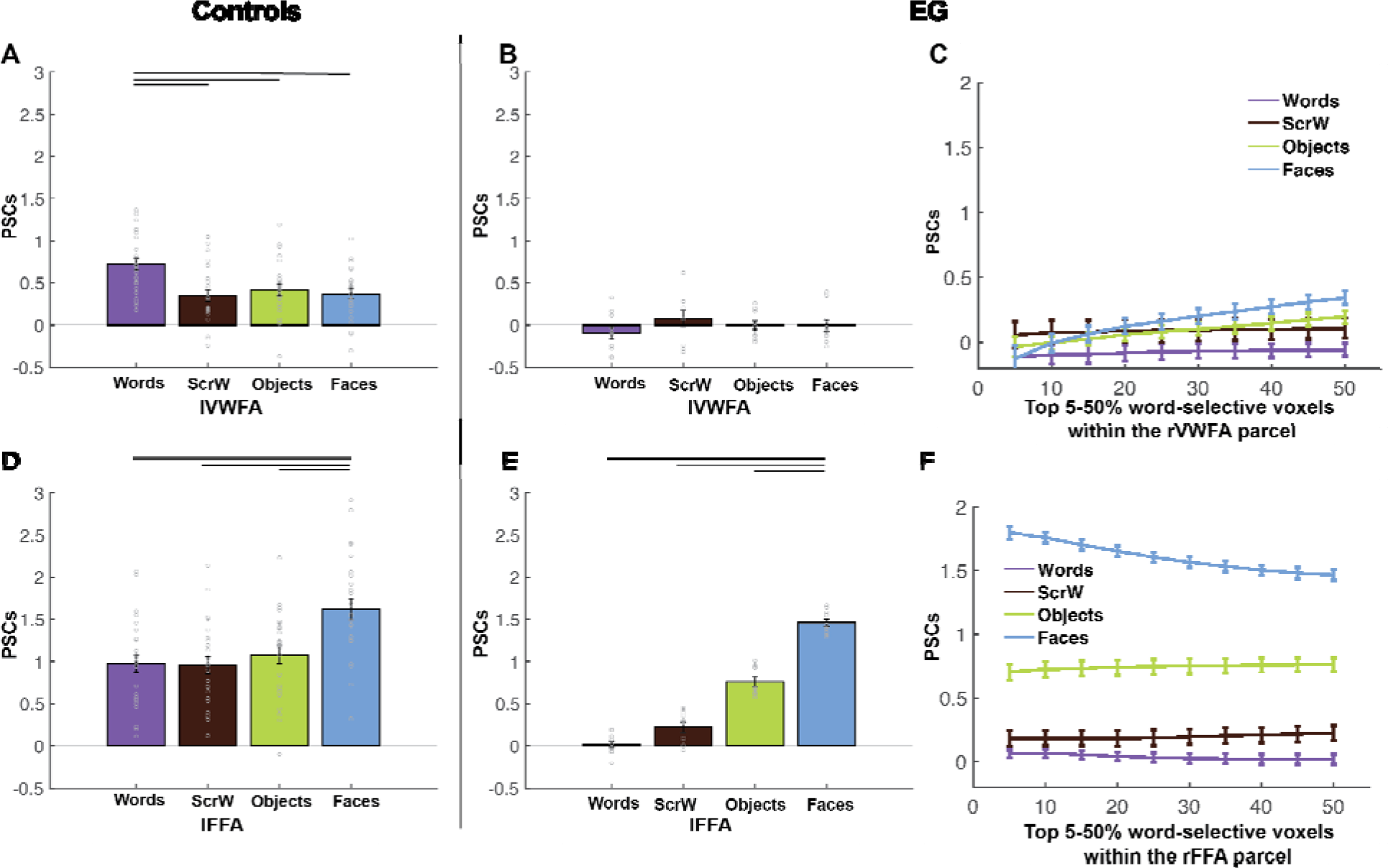
Responses to four conditions in the lVWFA and lFFA for EG and the controls group. (A), Bar plots show mean PSCs to four conditions estimated in independent data within individually defined lVWFA fROIs (i.e., top 10% word-selective voxels within lVWFA parcel) for the control group. (B), Mean PSCs to the four conditions estimated in independent data within individually defined lVWFA fROIs for EG. Here and in E, the results are averaged across run combinations.. (C), Parametrically decreasing the threshold for defining the lVWFA fROIs from the top 5% to 50% word-selective voxels within the lVWFA parcel. Number of voxels in the defined lVWFA fROIs: 5%=295 voxels, 50%=2402 voxels. Here and in F, average PSCs across run combinations are shown for each threshold. (D), Mean PSCs across participants to the four conditions estimated in independent data within individually defined lFFA fROIs for the control group. (E), Mean PSCs to four conditions estimated in independent data within individually defined lFFA fROIs for EG. (F), Parametrically decreasing the threshold for defining lFFA fROI from the top 5% to 50% face-selective voxels within the lFFA parcel. Number of voxels in the defined lFFA fROIs: 5%=18 voxels, 50%=185 voxels. In the bar plots, dots correspond to individual data points (controls: n=25 subjects; EG: n=10 run combinations, from ten iterations). Horizontal bars reflect significant paired t-tests p < 0.05. Error bars in both the bar and line plots denote standard errors of the mean by participants (for the control group) and by run combinations (for EG).

Similar to the analyses we performed for the rVTC, to test whether selectivity for other visual categories in the left VTC is typical in EG, we examined face selectivity in EG’s lFFA. EG’s face selectivity remained intact (**Figure 5E)** and did not differ from that of the control group (**Figure 5D**; t(24)=-0.244, p= 0.404, modified t-test; 95% Bayesian CI, [0.280 0.500]). EG’s lFFA fROI showed significantly higher responses to Faces than to other conditions (Faces vs. Words: t(9)=36.178, p= 4.663×10^-11^; Faces vs. Scrambled Words: t(9)=18.174, 2.109×10^-8^; Faces vs. Objects: t(9)=23.708, p= 2.016×10^-9^; **Figure 5E)**, just like what we observed in the control group (Faces vs. Words: t(24)=9.056, p=3.288×10^-9^; Faces vs. Scrambled Words: t(24)=8.865, p= 4.892×10^-9^; Faces vs. Objects: t(24)=6.880, p= 4.084×10^-7^; **Figure 5D**). As was the case for the rFFA, the face selectivity in EG’s lFFA was observed across all thresholds used to define the lFFA (**Figure 5F**). When examining the strength of selectivity, we found that controls showed significant (compared to zero) word selectivity in the lVWFA (t(24)= 6.093, p= 2.714×10^-6^) and face selectivity in the lFFA (t(24)= 6.569, p= 8.567×10^-7^). Interestingly, when we compared the strength of the selectivity, we found that EG’s word selectivity in the lVWFA was significantly lower than controls (t(24)= -1.767, p= 0.045, modified t-test; 95% Bayesian CI, [0.003 0.106]) but face selectivity in the lFFA was significantly higher than controls (t(24)= 2.305, p= 0.015, modified t-test; 95% Bayesian CI, [9.417×10^-5^ 0.046]).

Altogether, the examination of EG’s right and left VTC suggests that without the typical left-hemisphere frontotemporal language network from birth—and presumably without the necessary connections between these areas and parts of the VTC—a canonical VWFA, a word-selective area, does not develop in either hemisphere.

### Does the frontotemporal language network support visual word processing?

Finally, it is possible that while EG’s right or left VTC lacked word selectivity or linguistic selectivity, perhaps parts of her amodal frontotemporal language network show canonical univariate selectivity to visual words. We invited EG back and collected fMRI data for the VWFA localizer again but with whole-brain coverage (see Supplementary Methods for details). We first replicated our main results: we observed no canonical VWFA in the left or right VTC but found a typical FFA in both hemispheres (Supplementary Figure 3). Critically, whole-brain coverage allowed us to ask whether there is evidence of visual word selectivity in the frontotemporal language network.

Using the language parcels (Fedorenko et al., 2010) as our search space, we identified voxels that showed higher activation to Words > Others at different thresholds (from top 1% to top 10%).

Then in independent runs, we extracted activation to conditions in both the VWFA and visual and auditory language localizers. We failed to find any voxels in the frontotemporal language regions that showed Word-selective responses (Figure 6A-C): activation to visual Words was not differentiated from the other visual categories; activation for even auditorily presented stimuli were higher than the activation to visual Words (even though the fROIs were chosen with the visual Word contrast). We also identified language-selective fROIs by contrasting visually presented English sentences vs. Nonword sequences (e.g., Fedorenko et al., 2010; Tuckute et al., 2022); consistent with Tuckute et al. (2022), we also found that EG’s language fROIs was reorganized to the RH and showed significantly higher activation to English sentences than to Nonword sequences in both temporal and frontal cortex. But nonetheless, these language-selective fROIs showed selectivity to high-level linguistic information (regardless of visual or auditory modality) and did not show distinct activation to visual Words vs. other visual categories (Figure 6D-F) suggesting that these language fROIs are indeed selective to linguistic information rather than orthographic information.

**Figure 6.**
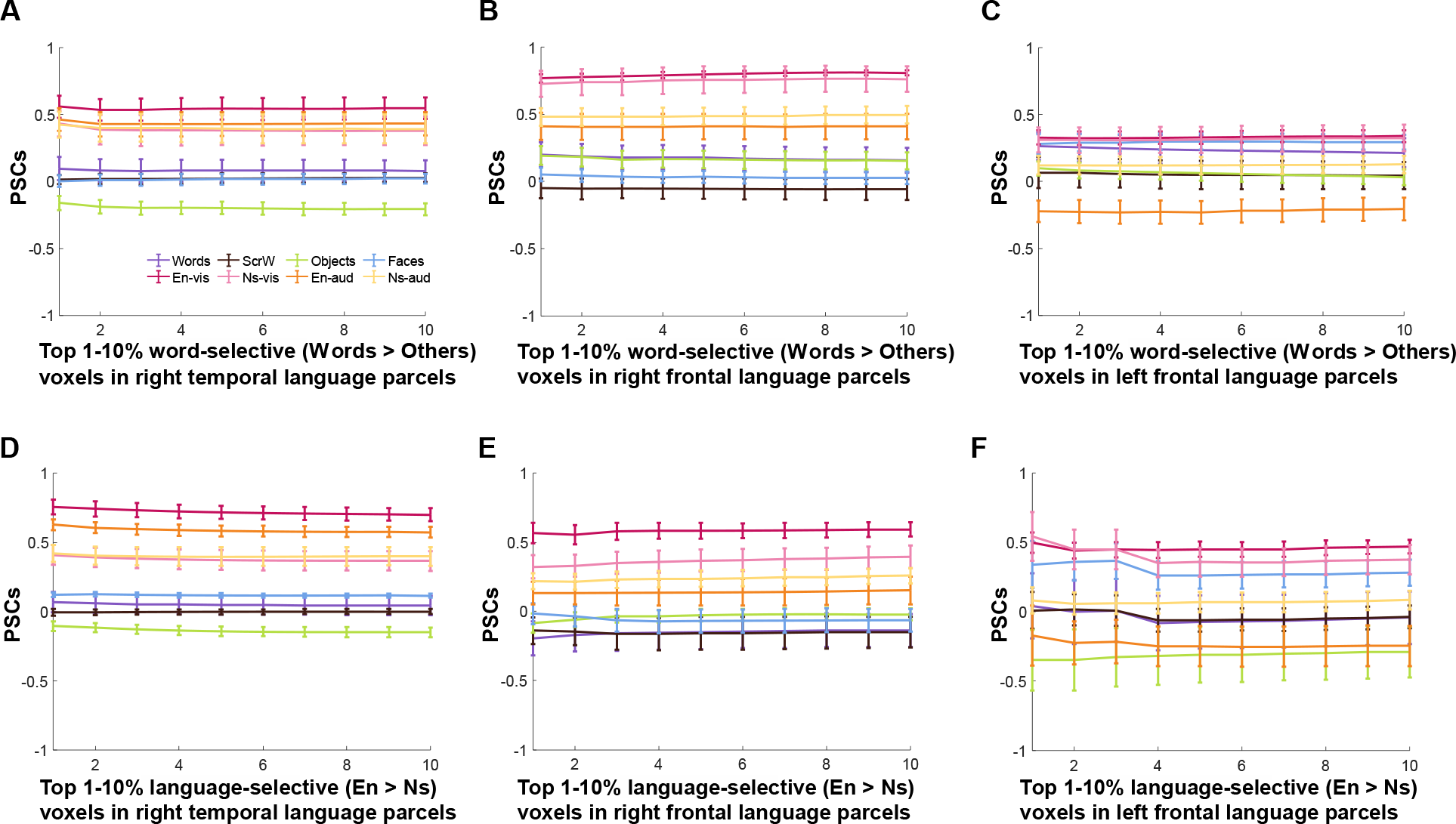
Responses to conditions in both the VWFA and language localizers in the frontotemporal language network for EG. (A-C), Mean PSCs in word-selective voxels (Words > Others) at different thresholds in the right temporal (A), right frontal (B) and left frontal language parcels (C). (D-F), Mean PSCs in language-selective voxels (Words > Others) at different thresholds in the right temporal (D), right frontal (E) and left frontal language parcels (F). Parametrically decreasing the threshold from the top 1% to 10% within each language parcel (i.e., the search space). Mean PSCs across run combinations (from 10 iterations for the VWFA task and 6 iterations for the language task) are shown for each threshold. Error bars denote standard errors of the mean by run combinations for EG. En-vis, visually presented English sentences; Ns-vis, visually presented Nonword sequences; En-aud, auditorily presented English sentences; Ns-aud, auditorily presented Nonword sequences.

### Distributed neural representation of visual words: multivariate pattern analysis (MVPA)

Our univariate analyses measured the average activation across the voxels most responsive to the visual category in question. Although this is the classic approach to demonstrate category selectivity, univariate analyses may be insensitive to potentially meaningful distributed representation patterns in suprathreshold and/or subthreshold voxels. Previous studies have found that categorical information can be reliably decoded by comparing within-category versus between-category correlations in the VTC (e.g., Haxby et al., 2001). Moreover, a recent study found mature representational similarity structures via multivariate patterns in children with no univariate selectivity, suggesting that distributed representations may developmentally precede category selectivity (Cohen et al., 2019). To explore whether words may be represented in a distributed fashion in EG, we performed a series of multivariate pattern analyses (MVPA).

We first examined the representational similarity matrices (RSMs) in the entire VTC (see Methods) to investigate whether multivariate representational structure for visual categories was preserved in EG. Indeed, we found that EG’s rVTC RSM was strongly and significantly correlated with that of the control group (r=0.918, p= 9.100×10^-3^ (permutation test); Supplementary Figure 5). We tested whether this correlation between EG and the control group was different from the correlations between any given control individual and the rest of the control group (see Methods). Single case comparisons showed that the RSM correlation for EG vs. controls did not significantly differ from the within-controls correlations (t(24)=0.797, p= 0.217, Bayesian 95% CI [0.09 0.351]). Similar results were found for the lVTC: the RSMs of EG and the control group were strongly and significantly correlated (r=0.888, p= 7.300×10^-3^), and the correlation for EG vs. controls did not significantly differ from the within-controls correlations (t(24)=0.835, p= 0.206, Bayesian 95% CI [0.09 0.34]).

We then asked whether EG’s ‘VWFA’ may contain voxels that show distinct distributed activation patterns to visual words. Specifically, as described in Methods, we searched for voxels that satisfied the following criteria: the searchlight around a given voxel should show 1) distinctive response patterns to Words vs. other categories (e.g., the Words-Words correlation should be higher than the Words-Faces correlation); and 2) stronger within-category correlations for the preferred category (e.g., the Words-Words correlation should be higher than the Faces- Faces, Objects-Objects, and Scrambled Words-Scrambled Words correlations). Previous studies have shown distributed representations within category-selective regions (e.g., the FFA) of non-preferred categories (e.g., places), and debate is ongoing over whether this information has functional relevance (e.g., Kanwisher, 2010). Our second criterion was included to identify voxels that show more stable multivariate representations for the category of interest (e.g., Words) compared to other categories. Indeed, we identified a set of voxels within the rVWFA and lVWFA parcels that showed a reliable distributed code for words in both controls and EG (Supplementary Figure 4A, 2C; Supplementary Table 4). In addition, mirroring the univariate analyses, we also identified a set of voxels that showed a reliable distributed code for faces within the rFFA and lFFA parcels in both controls and EG (Supplementary Figure 4B, 4D; Supplementary Table 4).

Critically, to test whether these distributed responses to visual words differ between EG and the controls, we subtracted the average within-category correlations for all non-word categories from the within-category correlations for Words. This difference tells us how much stronger the within-category correlation is for the preferred compared to the non-preferred categories. EG showed comparable within-category correlation differences to the controls in both the rVWFA (t(24)=1.081, p= 0.145, modified t-test, 95% Bayesian CI [0.05, 0.26] and lVWFA (t(24)=0.184, p= 0.428, modified t-test, 95% Bayesian CI [0.30, 0.50]); **Figure 7A**). Similar within-category correlation differences were also observed between EG and the controls in the rFFA (t(24)=- 0.802, p= 0.215; 95% Bayesian CI [0.09, 0.34]) and lFFA (t(24)=0.488, p= 0.315; 95% Bayesian CI [0.18, 0.46]); **Figure 7B**).

**Figure 7.**
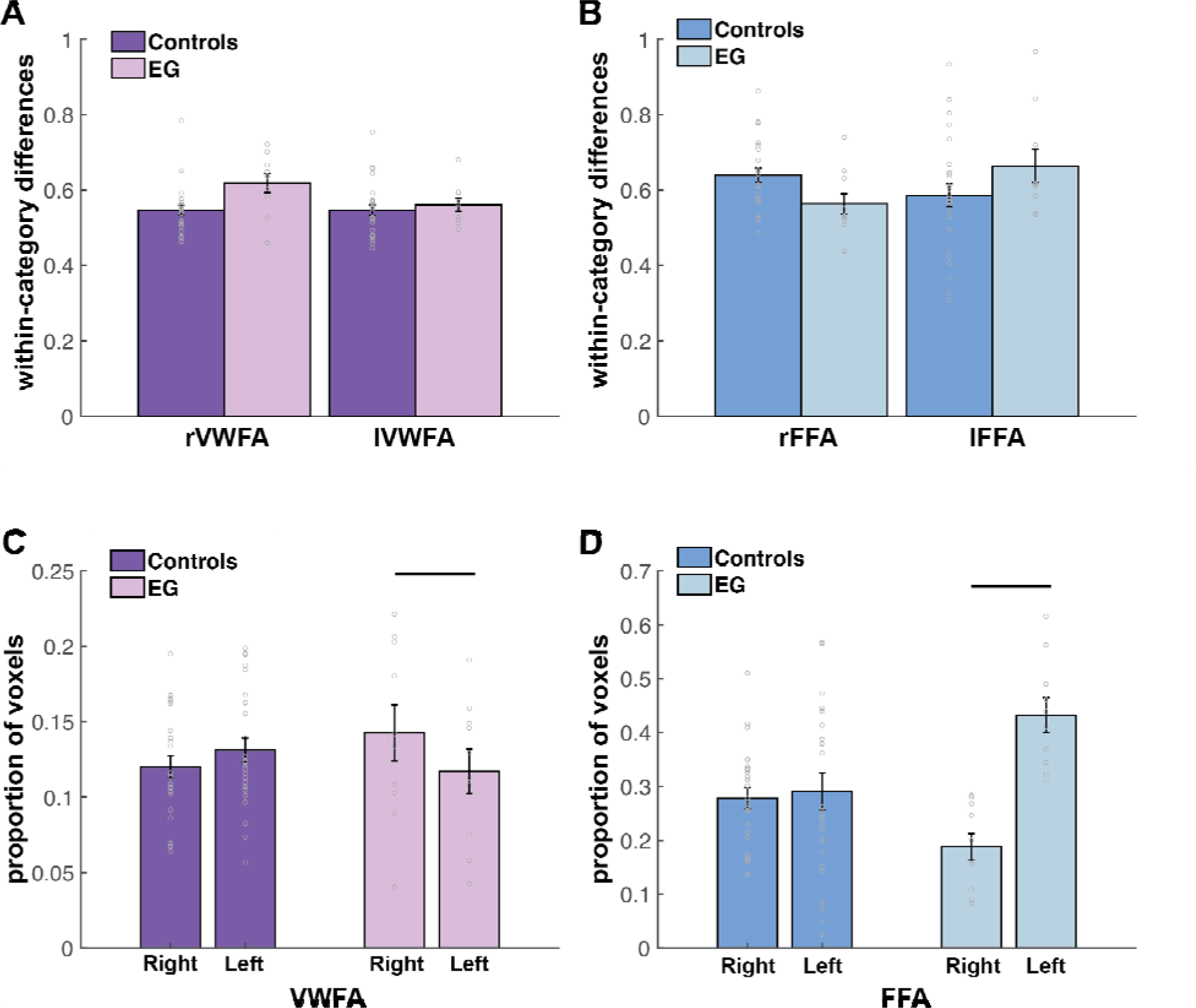
Results of the multivariate pattern analysis (MVPA). (A) Within-category correlation differences between preferred (i.e., Words-Words) and non-preferred (i.e., average within-category correlation of non-word conditions) conditions for EG and the controls in the rVWFA and lVWFA parcels. Here and in B, correlation values were Fisher’s z-transformed. (B) Within-category correlation differences between preferred (i.e., Faces-Faces) and non-preferred (i.e., average within-category correlation of non-face conditions) conditions for EG and the controls in the rFFA and lFFA parcels. (C) Proportion of voxels that show multivariate selectivity for Words in the rVWFA and lVWFA parcels for EG and the controls. (D) Proportion of voxels that show multivariate selectivity for Faces in the rFFA and lFFA parcel for EG and the controls. In the bar plots, dots correspond to individual data points (controls: n=25 subjects; EG: n=10 run combinations from ten iterations). Horizontal bars reflect significant LH-RH differences, p<0.05.Error bars denote standard errors of the mean by participants (for the control group) and by run combinations (for EG).

Finally, we explored potential hemispheric asymmetries with respect to multivariate selectivity for words and faces. Left-hemisphere dominance for words and right-hemisphere dominance for faces are well established with univariate measures. Do multivariate patterns also show these asymmetries? We calculated the number of voxels that show multivariate selectivity for words and faces within the left and right VWFA and FFA parcels, respectively (controlling for the size of the search space; see Methods). For Words, mirroring the univariate results in the past literature, a numerically larger proportion of voxels showing multivariate word selectivity was found in the left compared to the right VWFA in the controls (although the difference did not reach significance: t(24)=1.181, p=0.249). Interestingly, the opposite pattern was observed in EG, with a larger proportion of voxels showing multivariate word selectivity found in the right VWFA (t(9)= -2.469, p= 3.564×10^-2^) (**Figure 7C**). For Faces, a numerically larger proportion of voxels showing multivariate face selectivity was found in the right compared to the left FFA in the controls (although, like for Words, the difference did not reach significance: t(24)=0.367, p=0.717), and EG displayed the opposite pattern: a significantly larger proportion of voxels showing multivariate face selectivity in the left FFA (t(9)=8.983, p=8.672×10^-6^) (**Figure 7D)**.

## Discussion

Case studies provide valuable insights for understanding the functional organization of the human brain and the patterns of reorganization that follow neurological damage (Vaidya et al., 2019). In the current study, we had a unique opportunity to collect fMRI data from an individual (EG) born without her left superior temporal lobe in order to ask whether category-selective responses to visual words within the ventral temporal cortex would be affected. Specifically, we asked: in the absence of a typical left-hemisphere language network, does word selectivity emerge, and if so, where? Does the VWFA emerge in the right VTC given that EG’s language network is located in the right hemisphere (Tuckute et al., 2022)? Or do we instead, or additionally, observe word selectivity in the canonical VWFA within the spared lVTC? Surprisingly, we found no univariate word selectivity within the rVWFA, the right homotope of the canonical lVWFA, and even in an expanded search space of the entire rVTC. We also did not observe word-selective responses in the lVWFA or in any other spared parts of the lVTC. Moreover, the response magnitude to Words (relative to baseline) was significantly lower in EG’s lVWFA than that of controls. Importantly, this lack of category selectivity was specific to Words: selective responses to Faces remained intact in EG’s rFFA and lFFA.

Absent univariate word selectivity, we then explored multivariate representations of visual words in EG. We found that EG’s VTC showed an overall similar representational structure to that of the control group, and that, like the control group, EG had a set of voxels whose local neighborhoods robustly differentiated between Words and other visual categories in their patterns of response. Critically, these voxels also showed a higher within-category correlation (across runs) for Words compared to the within-category correlations for other categories, and the degree of this ‘multivariate selectivity’ was similar between EG and the controls.

Interestingly, however, in contrast to the typically observed left-hemispheric dominance for words and right-hemispheric dominance for faces, EG had a larger proportion of voxels that showed multivariate selectivity for words in her right than her left VWFA, and a larger proportion of voxels that showed multivariate selectivity for faces in her left FFA than her right FFA.

Altogether, the current study suggests that in the absence of a typical left-lateralized language network—at least in cases of early left-hemispheric damage, when language processing has no choice but to localizer to the right hemisphere—neural selectivity for visual words is, or at least can be, highly atypical. In our participant with extensive left-hemispheric temporal damage and a right-lateralized language network, we observed no word selectivity in the univariate analyses in either the right or the spared left VTC, replicated across scanning sessions (5 years apart). The absence of such selectivity, combined with EG’s intact reading ability, suggests that successful orthographic processing may depend on a more distributed and more right-lateralized neural representation than that observed in typical literate adults.

### Canonical univariate word selectivity may not emerge when a left-hemisphere language network cannot develop normally

The interesting case of EG allowed us to investigate how visual word processing within the VTC can be affected by a congenital or early left-hemisphere lesion outside of visual cortex. Our results provide the first evidence of atypical visual word selectivity in the VTC when the left-lateralized language cortex is missing, and when the language network consequently develops in the right hemisphere (Tuckute et al., 2022) during the early stages of language learning and prior to learning to read. Our study also suggests that even with some remaining anatomical connections between the spared lVTC and the frontal and temporal areas (presumably via local U-fibers or remaining long-range fibers), the lVWFA will not emerge at the stereotypical location when the left hemisphere does not support language processing.

How does a left language network contribute to the development of the VWFA? One possibility is that in a typical brain, the site of the putative VWFA is predisposed to written scripts via intrinsic co-activation with speech and/or language/semantic areas in the left temporal (and maybe frontal) cortex. The processing of speech sounds is tightly linked to reading, and impaired coding of speech sounds (e.g., phonological awareness) is often considered a precursor of dyslexia (e.g., Pennington & Bishop, 2008; Shaywitz et al., 2002). In the absence of regions that typically support speech processing (within the left superior and middle temporal gyri (STG and MTG); Raschle et al., 2012), EG’s left VTC lacked early interactions with these regions. Further, a typical lVWFA communicates visual orthographic information to higher-level left temporal (and maybe frontal) regions that integrate visual and auditory information, like the amodal language regions that process lexical-semantic and combinatorial linguistic information (e.g., Fedorenko et al., 2020) and the regions that support abstract conceptual processing (e.g., Binder et al., 2009; Patterson et al., 2007)—both sets of areas that were also missing in EG’s left hemisphere. Consistent with the idea that connections between the left VTC and the ipsilateral high-level language areas may be critical for the emergence of the lVWFA, a recent study found that in newborns, the putative site of the lVWFA already shows preferential functional connectivity to the areas that will later respond to high-level linguistic information, compared to adjacent regions (Li et al., 2020); this pre-existing functional coupling may further strengthen during language and reading acquisition. Because EG was missing both i) speech-responsive areas, and ii) higher-level language/conceptual areas in her left hemisphere, we cannot evaluate the relative contributions of these two sets of areas and their connections with the VTC to the emergence of a canonical VWFA. We speculate that both may be important.

### Right-hemispheric neural correlates that support visual word processing

Because EG’s language network resides in the right hemisphere (Tuckute et al., 2022), we expected to find a VWFA in the right VTC. Surprisingly, no univariate word selectivity was observed in EG’s VTC, and this result could not be explained by either a lack of functional specialization *anywhere* in the rVTC (intact face selectivity was observed), or our particular procedure for defining the VWFA (the results held across a range of thresholds, when using a larger search space, and across different contrasts). Interestingly, neurotypical individuals with right-dominant language activation sometimes also lack a VWFA in the right VTC (e.g., Cai et al., 2008; Van der Haegen et al., 2012); in these individuals, lVTC appears to be engaged during word recognition, presumably due to stronger frontotemporal anatomical connections in the LH than in the RH, and any language activation on the left (even if it’s non-dominant) would engage the lVTC for reading (Powell et al., 2006). In the case of EG, her right-hemisphere language network may have lacked early interactions with the rVTC, and her left-hemisphere language network was altogether lacking, resulting in the atypical word selectivity (in both right and left) that we observed here.

Some have also argued that the development of word-selective cortex directly competes with face-selective cortex for neural resources, thus contributing to right-hemispheric dominance for face processing and left-hemispheric dominance for word processing in neurotypical individuals (e.g., Behrmann & Plaut, 2015; Dehaene & Cohen, 2007). Interestingly, EG showed typical face selectivity in the right FFA, not different from controls. It is therefore possible that a focal word-selective area failed to emerge in EG’s rVTC because the relevant cortical tissue had already been ‘assigned’ to face processing, perhaps due to some specific visual features of faces that are better represented in the right hemisphere/left visual field (e.g., holistic face processing; Li et al., 2017; Rossion et al., 2000). Interestingly, although the strength of the face selectivity did not differ between EG and controls in the rFFA, EG showed significantly higher face selectivity in the lFFA. In line with this finding, more multivariate-selective face voxels were present in EG’s lFFA, compared to her rFFA, in sharp contrast to controls. The functional significance of the latter is at present unclear, but can be explored in future work. On the other hand, EG had more multivariate-selective word voxels in her rVWFA than her lVWFA, presumably related to the fact that her language network resides in the right hemisphere. These data suggest that the hemispheric dominance of the language network drives the laterality of visual processing in the VTC (be it implemented focally, or in a distributed fashion), at least for words, but perhaps also for faces (Behrmann & Plaut, 2020; Dehaene & Cohen, 2007).

Further, we found no evidence in support of theories that propose that the VWFA does not process orthography (despite evidence in our neurotypical control population) and that it instead processes linguistic stimuli in much the same way as the lateral frontotemporal amodal language network (see Price & Develin 2011). We investigated multiple ways of defining Word or Language-selective fROIs in the right and left VTC, and found no evidence that EG’s VTC is selective to either general linguistic stimuli or visual Word stimuli despite normal reading performance.

### No word-selective response observed outside the VTC

Although we show that normal reading ability is possible in the absence of focal selectivity for word processing in the VTC, there may be other pathways or neural structures outside of the VTC that are important for EG’s reading ability. For example, Seghier et al. (2012) reported a patient who acquired dyslexia following extensive left ventral occipitotemporal cortex (LvOT) resection, but then regained reading ability; they provided evidence to suggest that the patient’s reading ability was now supported via a direct connection between the occipital visual cortex and the left superior temporal sulcus (STS) without the involvement of the ventral visual stream. It is therefore possible that the STS in the language-dominant hemisphere (i.e., the right hemisphere in EG) may support visual word processing through its connections with the occipital cortex.

Thus, to supplement our main analyses, which were restricted to the ventral temporal cortex, we invited EG back and obtain the fMRI data from the same VWFA task with a whole-brain coverage. We found no evidence of significantly greater responses to visual Words as compared to other visual stimuli in the lateral temporal or frontal cortex, either when searching for word-selective voxels directly (by Words > Others) or looking for word selectivity in language fROIs (canonically defined with English sentences > Nonword sequences). These results further confirmed the lack of univariate Word-selective responses in EG.

### Multivariate responses to words

Despite the absence of typical univariate word selectivity, EG’s VTC was similar in its overall representation similarity structure—across different visual stimulus classes—to that of the control group. Consistent with our results, Liu et al. (2019) also found typical representational structure in category-selective regions after resections within or outside VTC; in addition, a recent study found mature representational similarity structures in children (5-7 year-olds) with no univariate selectivity, suggesting that distributed representations precede category selectivity (M. A. Cohen et al., 2019). Our results provide another case where a typical multivariate representational structure is observed in the absence of univariate selectivity for words. Moreover, our results showed that EG (like typical adults) displayed distinctive activation patterns within the general vicinity of both right and left VWFA that could distinguish visual words from other conditions, with higher within- than between-category correlations (Haxby et al., 2001) and more robust (consistent over time) representations of Words than other conditions. Supporting the idea that multivariate selectivity for words may be functionally useful, Stevens et al. (2017) found the VWFA (defined in individual participants using a standard univariate contrast) discriminated words from pseudowords, and did so more strongly than other control regions (e.g., the FFA).

It remains unknown that what the relationship is between univariate and multivariate representations. Univariate selectivity may partially depend on a connectivity scaffold or other genetically defined instructions to determine the location of functional specialized areas (Deen et al., 2017; Kamps et al., 2020; Li et al., 2020). In contrast, distributed representations like those found in EG may rely less on a connectivity scaffold. Do distributed representations reflect a fundamentally different neural code from that associated with focal representations? Some have argued that the answer is no, and that multivariate analyses may simply be more sensitive than univariate approaches given that they consider the heterogeneity of response across voxels within a region as well as potential subthreshold voxels (Davis et al., 2014). On the other hand, multivariate representations may be truly distinct and reflect information derived from bottom-up visual statistics to a greater extent; and thus perhaps in typical development, representational similarity structures precede univariate selectivity (M. A. Cohen et al., 2019) and emergence of univariate selectivity requires experience-dependent interactions with higher-level areas (e.g., the language cortex in case of the VWFA). Speculating further, perhaps during EG’s early stages of language development (while her language system was maturing in the right hemisphere), the right visual cortex lacked the critical early interaction with the language cortex, due to the lack of strong innate connectivity between the two in the right hemisphere (cf. the left hemisphere; e.g., Li et al., 2020; Powell et al., 2016), resulting in a more distributed representation for visual words.

### Limitations and future directions

Overall, our study provides unique evidence that in the absence of a typical left-hemisphere language network—at least in cases of early left-hemisphere damage—the canonical word-selective lVWFA may not develop. Some limitations are worth noting. First, our results raise an interesting question about the behavioral relevance of category-selective regions. Previous studies have found that the strength of univariate category-selective selectivity (or lack thereof) accounted for performance differences in various object recognition tasks (e.g., Furl et al., 2011; Grill-Spector et al., 2004; Huang et al., 2014; Song et al., 2015). The VWFA exists due to reading and literacy and is not observed in illiterate individuals (e.g., Dehaene et al., 2015; Saygin et al., 2016) and disrupted activity in the VWFA is observed in dyslexic individuals (e.g., Centanni et al. 2019; Maisog et al., 2008; Paulesu et al., 2014; Richlan et al., 2011). Additionally, lesions or disruption of the VWFA with brain stimulation, results in failures in word recognition (Hirshorn et al., 2016). All this evidence underscores the behavioral importance of the VWFA in reading. The unique case of EG, who has normal reading ability without univariate word selectivity, suggests an alternative mechanism that might support orthographic processing. Here we examined one possibility that visual word processing might be represented in a distributed manner. Future work can further investigate this possibility. For example, it would be interesting to explore how multivariate representations are read out by downstream brain regions for further processing (Williams et al., 2007), or to further test the link between behavioral performance and multivariate representations. A previous study showed that performance in a visual search task among different pairs of object categories could be predicted from underlying neural similarity structures (M. A. Cohen et al., 2016). At this point, it is unclear how multivariate representations for words contribute to EG’s reading ability, and whether the right-hemisphere representations are more important than the left-hemisphere ones—questions that will be a focus of future studies. Second, EG’s missing left superior temporal cortex led to right-lateralized speech and language processing. As discussed earlier, it remains unclear whether speech and higher-level language areas contribute equally to the emergence of the lVWFA, and whether temporal language regions may be more important than the frontal ones. Finally, here we reported a single case that sheds new light on the role of speech and language areas in developing a VWFA. Generalization to other, similar cases, and data from other methods like noninvasive brain stimulation will help us better understand the causal role of language cortex in developing visual word selectivity.

## Acknowledgements

We would like to thank our participant – EG – for her time and willingness to participate in brain research. We thank Greta Tuckute, Alexander Paunov, and Benjamin Lipkin for their help in coordinating the collaboration as well as their suggestions for data analysis. We also thank Idan Blank and Zachary Mineroff for help with EG’s fMRI and behavioral data collection, Rachel Ryskin for sharing the data from the speeded reading task, and members of Saygin Developmental Cognitive Neuroscience Lab for helping with control data collection. We would like to acknowledge the Athinoula A. Martinos Imaging Center at the McGovern Institute for Brain Research, MIT and the support from Center for Cognitive and Behavioral Brain Imaging (CCBBI) and Ohio Supercomputer Center (OSC). E.F. was supported by NIH awards R01 DC016607, R01 DC016950, and research funds from the McGovern Institute for Brain Research, the Department of Brain and Cognitive Sciences, and the Simons Center for the Social Brain at MIT. Z.M.S. was supported by the Alfred P. Sloan Foundation and OSU’s Arts & Sciences College and Chronic Brain Injury initiative.

## Conflict of interest

The authors declare no competing financial interests.

## Notes

### Competing Interest Statement

The authors have declared no competing interest.

