## Supplementary Information for "Intact reading ability in spite of a spatially distributed visual word form ‘area’ in an individual born without the left superior temporal lobe"

Supplementary Methods: Data acquisition for the VWFA localizer with a whole-brain coverage

Supplementary Table 1: Comparing EG’s reading ability with neurotypical individuals

Supplementary Table 2: Comparison between results in a small group of adults scanned at MIT to our control group (adults scanned at OSU from main text)

Supplementary Table 3: Reading component (task #18-38) of PALPA (Kay et al., 1992)

Supplementary Figure 1. Searching word-selective voxels within the VTC for EG

Supplementary Figure 2. Activation to a visual-audio language task within the VTC for EG

Supplementary Figure 3. Responses to four conditions in the bilateral VWFA and FFA fROIs for EG (Replication of the main results using data from EG’s recent visit)

Supplementary Figure 4: Split-half within-category correlations to four conditions in EG and the control group.

Supplementary Table 4: Comparing split-half within-category correlations for words and faces between EG and the control group

Supplementary Figure 5: Representation similarity matrices (RSMs) of the VTC in EG and the control group

**Supplementary Methods: Data acquisition for the VWFA localizer with a whole-brain** **coverage**

To explore potential VWFA reorganization outside the VTC and in the right temporal and/or frontal cortex, we invited EG back in November 2021. EG completed four runs of the same VWFA localizer but with a whole-brain coverage. Specifically, functional images were acquired with the echo-planar imaging (EPI) sequence: TR = 2000ms, TE = 30ms, 172 TRs, 100 × 100 base resolution, voxel resolution = 2.2 mm^3^, field of view (FOV) = 220mm, 54 slices acquired in an interleaved order to cover the whole brain.

Supplementary Table 1: Comparing EG’s reading ability with neurotypical individuals

1a: Comparing EG’s performance during the VWFA localizer with the current control group (Crawford & Howell's modified t-test).

| 1-back VWFA localizer task (EG vs. current controls (N=25), Crawford-Howell modified t-test) | | | | |
| --- | --- | --- | --- | --- |
| Accuracy | | | | |
|  | Words | ScrW | Objects | Faces |
| p | 0.311 | 0.500 | 0.392 | 0.425 |
| t | 0.500 | 0.000 | -0.278 | -0.191 |
| df | 24 | 24 | 24 | 24 |
| Response Time | |  |  |  |
|  | Words | ScrW | Objects | Faces |
| p | 0.334 | 0.336 | 0.110 | 0.387 |
| t | 0.434 | 0.430 | 1.257 | -0.290 |
| df | 24 | 24 | 24 | 24 |

1b: Comparing EG’s performance in a speed-reading comprehension task with a large sample of neurotypical adults (Crawford & Howell's modified t-test).

| Speed-reading comprehension task (EG vs. Normatives (N=191), Crawford-Howell modified t-test) | | | | | | | | |
| --- | --- | --- | --- | --- | --- | --- | --- | --- |
| Speed | 1 | 2 | 3 | 4 | 5 | 6 | 7 | 8 |
| p | 0.390 | 0.494 | 0.461 | 0.440 | 0.374 | 0.363 | 0.286 | 0.415 |
| t | 0.279 | 0.016 | -0.098 | -0.151 | 0.321 | -0.350 | -0.565 | 0.216 |
| df | 190 | 190 | 190 | 190 | 190 | 190 | 190 | 190 |

Supplementary Table 2: Comparison between results in a small group of adults scanned at MIT to our control group (adults scanned at OSU from main text)

| **PSCs in lVWFA** | | |
| --- | --- | --- |
|  | OSU vs. MIT^a^ | EG vs. MIT^b^ |
| Words | t(35)= 0.899, p= 0.375 | **t(11)= -2.353, p= 0.019** |
| SrwW | t(35)= 0.660, p= 0.514 | t(11)= -0.878, p= 0.199 |
| Objects | t(35)= -0.110, p= 0.913 | t(11)= -0.955, p= 0.180 |
| Faces | t(35)= 0.818, p=0.419 | t(11)= -1.173, p= 0.133 |
| **Selectivity** | | |
|  | OSU vs. MIT | EG vs. MIT |
| Words (lVWFA) | t(35)= 1.471, p= 0.150 | **t(11)= -1.613, p= 0.068** |
| Faces  (lFFA) | **t(37)= 2.098, p= 0.043** | t(13)= 0.440, p= 0.334 |

a, Two-sample t-test

b, Crawford-Howell modified t-test.

Comparable PSCs and selectivity levels were observed between MIT and OSU data within the lVWFA and lFFA; compared to controls scanned at MIT, consistent with main results, EG’s lVWFA showed significantly lower activation to words but not other conditions.

Supplementary Table 3: Reading component (task #18 through #38) of PALPA (Kay et al., 1992)

| **Task Number** | **Content** |
| --- | --- |
| 18 | mirror reversal (distinguishing between correct and mirror-reversed forms of individual letters); |
| 19 | upper case – lower case letter matching (finding the correct lower case form to match a given upper case letter); |
| 20 | lower case – upper case letter matching; |
| 21 | letter discrimination within words and nonwords (deciding whether pairs of items are the same, as in bribe-BRIBE, or different, as in BRIBE-tribe); |
| 22 | letter naming and sounding (naming or sounding the letters of the alphabet); |
| 23 | spoken letter – written letter matching (matching a heard letter with its appropriate written form); |
| 24 | visual lexical decision with ‘illegal’ nonwords (deciding whether a letter string is a word or not); |
| 25 | imageability and frequency visual lexical decision (deciding whether a letter string is a word or not, with the words varying in imageability and frequency); |
| 26 | visual lexical decision and morphology (deciding whether a letter string is a word or not; all letter strings consist of a root morpheme and a suffix morpheme, and the nonwords are made up of real morphemes, as in fullen or wises); |
| 27 | visual lexical decision and spelling-sound regularity (deciding whether a letter string is a word or not; half of the real words have irregular spelling, and half of the nonwords are homophonous with real words); |
| 28 | homophone decision (deciding whether two written words or nonwords are pronounced the same; e.g., bury-berry vs. fury-ferry, or quib-kwib vs. thib-shib); |
| 29 | letter length reading (out-loud reading of monosyllabic words varying in length from 3 to 6 letters); |
| 30 | syllable length reading (out-loud reading of 5-letter-long words varying in length from 1 to 3 syllables); |
| 31 | imageability and frequency reading (out-loud reading of words varying in imageability and frequency); |
| 32 | grammatical class reading (out-loud reading of words of different parts of speech: nouns, verbs, adjectives, and function words); |
| 33 | grammatical class x imageability reading (out-loud reading of nouns and function words that are similarly imageable); |
| 34 | lexical morphology and reading (out-loud reading of morphologically complex words: words with regular inflections, words with derivational endings, and irregularly inflected words); |
| 35 | spelling-sound regularity and reading (out-loud reading of regular and irregular words matched for frequency, imageability, part of speech, and length in letters, syllables, and morphemes); |
| 36 | nonword reading (out-loud reading of monosyllabic nonwords varying in length from 3 to 6 letters); |
| 37 | sentence reading (out-loud reading of sentences); |
| 38 | homophone definition and regularity (defining words from written forms alone; all words have a homophone with a different meaning; e.g., mail or suite) |

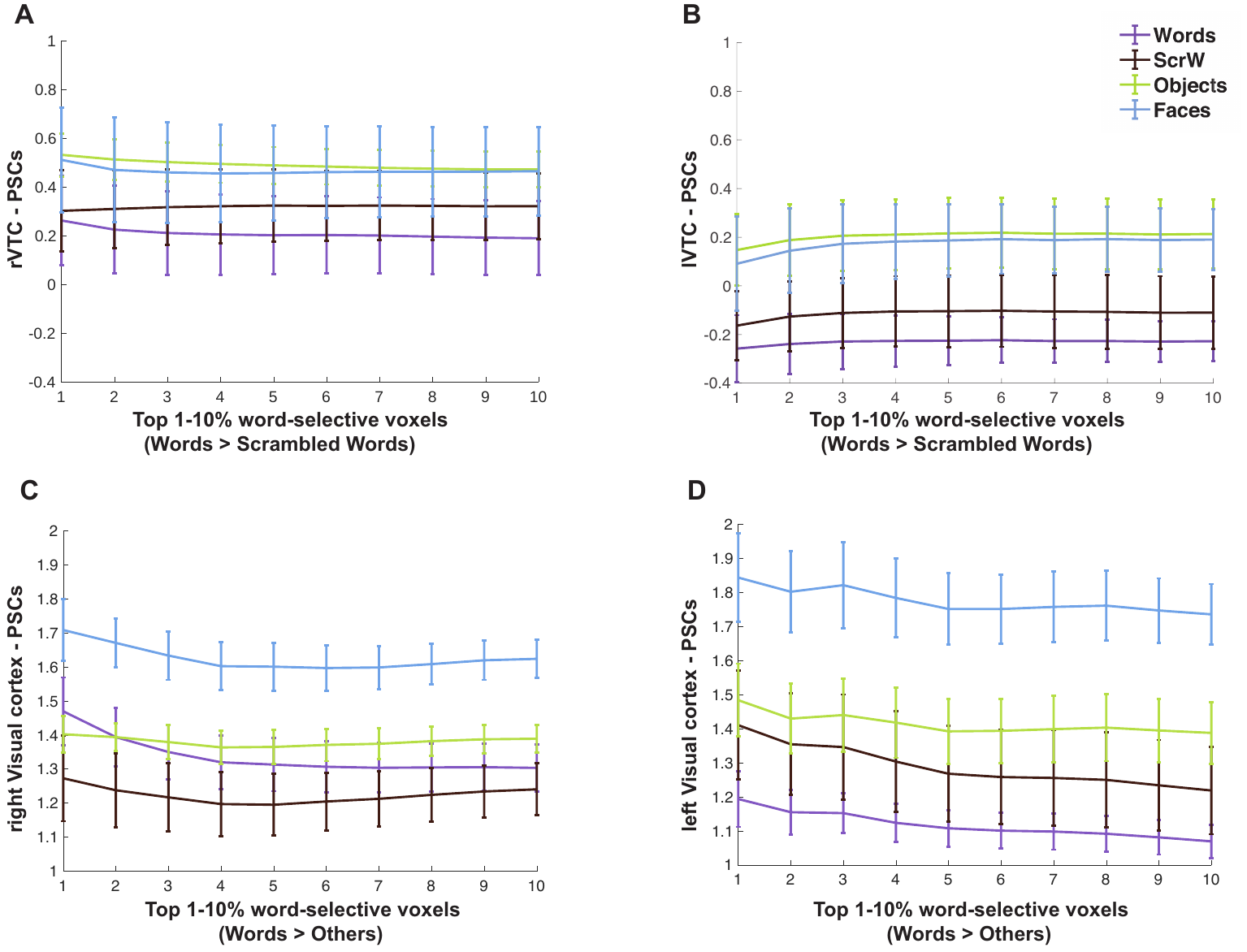

Supplementary Figure 1. Searching word-selective voxels within the VTC for EG. (A-B) Mean PSCs in word-selective voxels (Words > Scrambled Words) at different thresholds in the right and left VTC. (C-D) Mean PSCs in word-selective voxels (Words > Others) at different thresholds in the left and right visual cortex. A statistical map was created by contrasting activation to all four visual categories to the fixation block (All > Fix). We selected voxels that showed significant activation (p < 0.01) to at least half of the runs (3 out of 5 runs) as the visual mask for each hemisphere. Parametrically decreasing the threshold from the top 1% to 10% within the masks. Mean PSCs across run combinations (from 10 iterations) are shown for each threshold. Error bars denote standard errors of the mean by run combinations for EG. **This result showed that even with a less stringent contrast (Words > Scrambled Words) or a more restricted searching space (voxels that consistently involved in visual processing) and multiple thresholds to define word-selective voxels, no univariate response was found.**

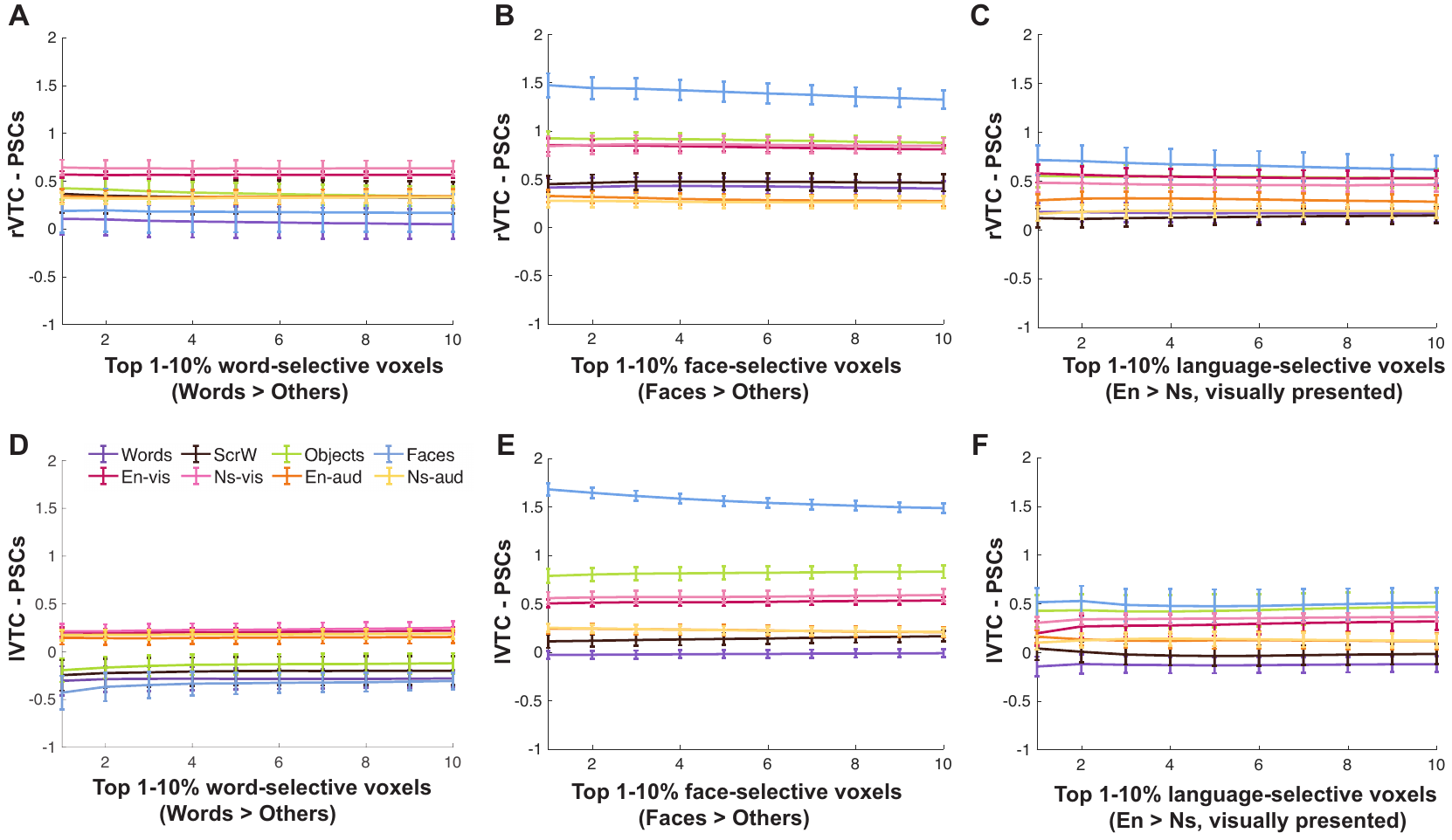

Supplementary Figure 2. Activation to a visual-audio language task within the VTC for EG. English sentences (En) and nonsense sentences (Ns) were presented either visually or auditorily during the task. (A) and (D), Mean PSCs in word-selective voxels (Words > Others) at different thresholds in the rVTC and lVTC. (B) and (E), Mean PSCs in face-selective voxels (Faces > Others) at different thresholds in the rVTC and lVTC. Activation to four visual categories also plotted as in the Figure 4 in the main manuscript. (C) and (F), Mean PSCs to conditions in both language and VWFA tasks were plotted for voxels that showed highest activation to visually presented English sentences (En > Ns) within the right and left VTC. Parametrically decreasing the threshold from the top 1% to 10% within the bilateral VTC. Mean PSCs across run combinations (from 10 iterations for the VWFA task and 6 iterations for the language task) are shown for each threshold. Error bars denote standard errors of the mean by run combinations for EG. **This result showed that there is no language selectivity in the VTC that would also support visual word processing.**

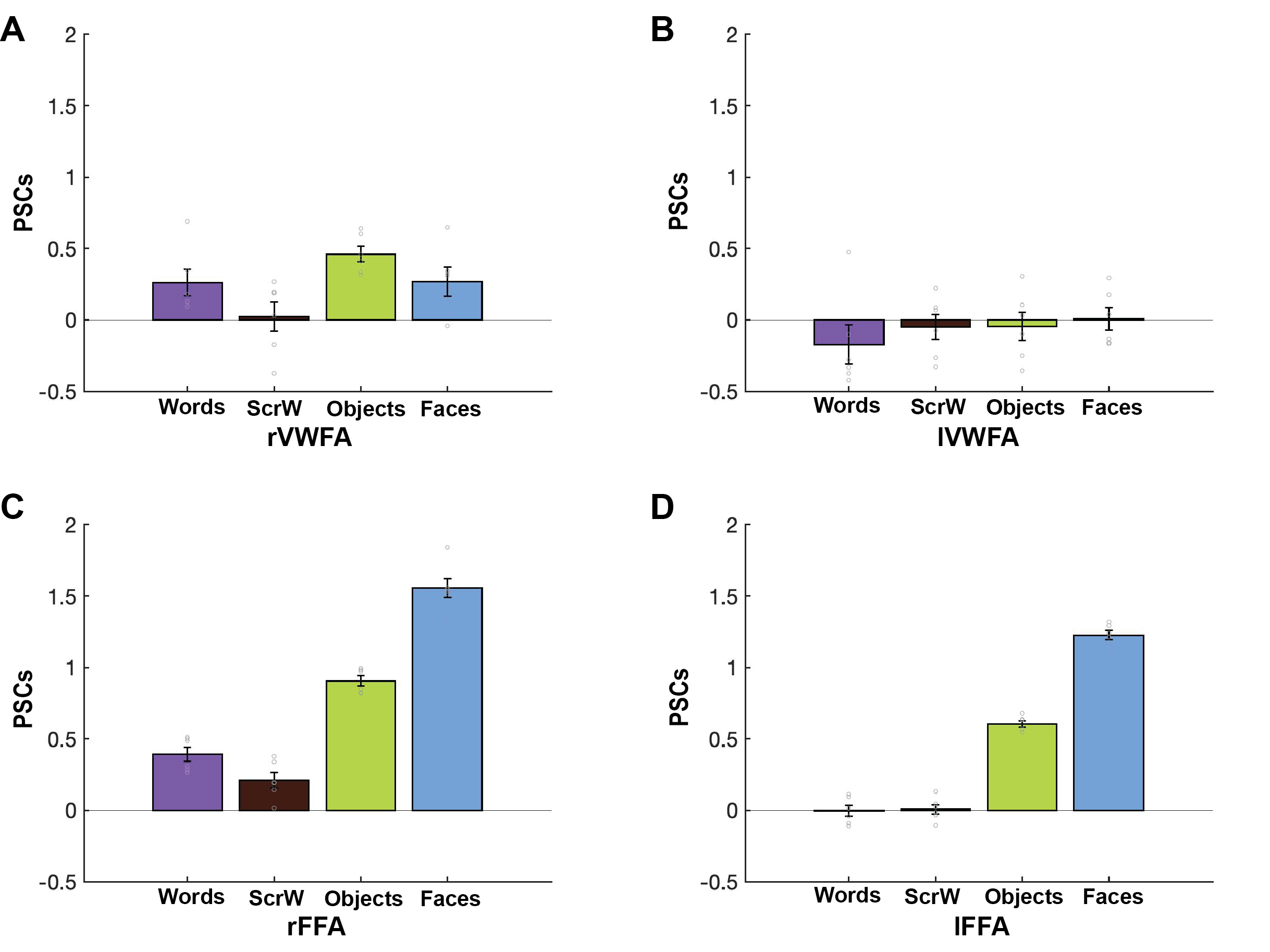

Supplementary Figure 3. Replication of the main results with EG’s data from a separate scanning session (5 years apart). (A-B), Mean PSCs to the four conditions estimated in independent data within the individually defined rVWFA (A) and lVWFA (B) fROIs for EG. (C-D), Mean PSCs to the four conditions estimated in independent data within the individually defined rFFA (C) and lFFA (D) fROIs for EG. The results are averaged across six run combinations. Error bars denote standard errors of the mean across run combinations.

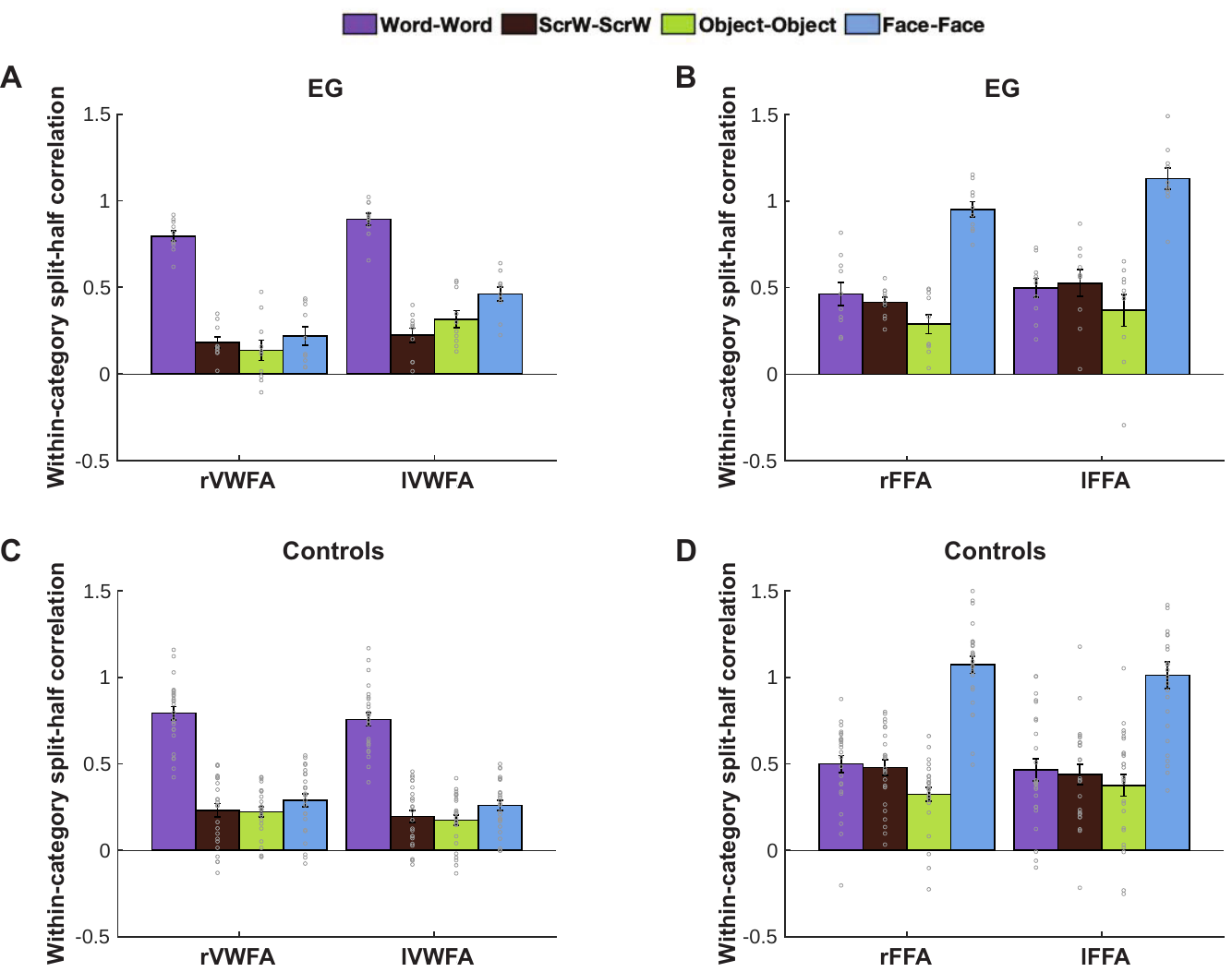

Supplementary Figure 4. Split-half within-category correlations to four conditions in EG and the control group. (A), Mean correlations were shown for voxels that represented word selectively within rVWFA and lVWFA parcels for EG. Here and in B, the results are averaged across run combinations. (B), Mean correlations were shown for voxels that represented faces selectively within rFFA and lFFA parcels for EG. (C), Mean correlations across participants were shown for voxels that represented words selectively within rVWFA and lVWFA parcels for controls. (D), Mean correlations across participants were shown for voxels that represented faces selectively within rFFA and lFFA parcels for controls. Correlation values were Fisher’s z-transformed. **Voxels that represented a given condition were chosen if they showed higher within- vs. between-category correlations for that condition and showed higher within-category correlation for that condition vs. within-category correlations of other conditions.**

Supplementary Table 4: Comparing split-half within-category correlations for words and faces between EG and the control group

| EG vs. Controls (Crawford & Howell's modified t-test) | | | | |
| --- | --- | --- | --- | --- |
|  | Words-Words | | Faces-Faces |  |
|  | lvwfa | rvwfa | lffa | rffa |
| p | 0.245 | 0.493 | 0.3841 | 0.3181 |
| t | 0.702 | 0.018 | 0.298 | -0.479 |
| df | 24 | 24 | 24 | 24 |

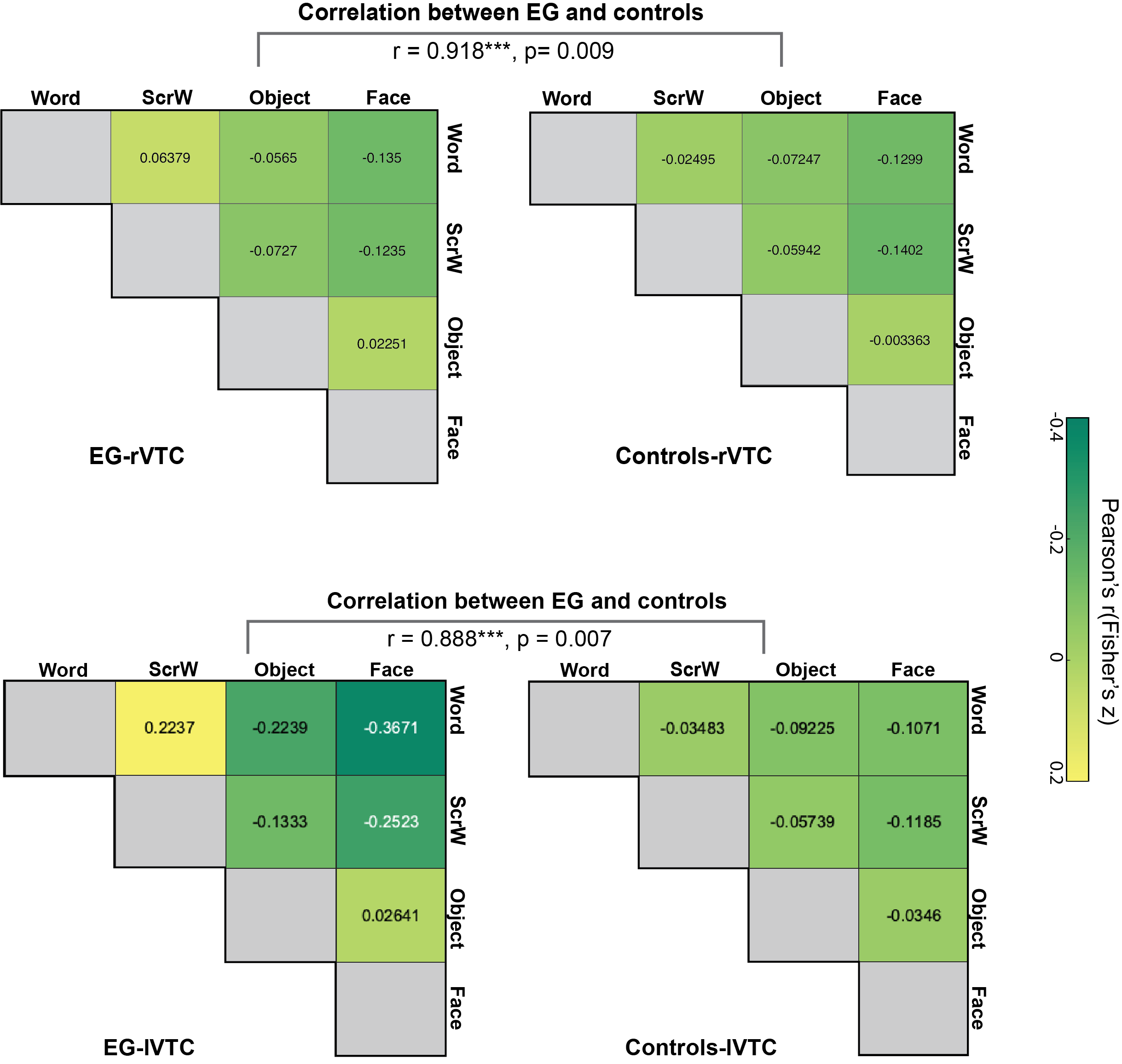

Supplementary Figure 5: Representation similarity matrices (RSMs) of the VTC in EG and the control group. RSMs were calculated for pairwise similarities of different conditions based on spatial response patterns of each searchlight; resulting correlation values (Fisher’s z-transformed) were averaged across all voxels within rVTC (Top) and lVTC (Bottom) for EG (Left) and the controls (Right) respectively. **This result showed that in both rVTC and lVTC, RSMs of EG were highly correlated with RSMs of the controls.**
